# Interferon ε restricts Zika virus infection in the female reproductive tract

**DOI:** 10.1101/2023.04.06.535968

**Authors:** Chuan Xu, Annie Wang, Laith Ebraham, Liam Sullivan, Carley Tasker, Vanessa Pizutelli, Jennifer Couret, Cyril Hernandez, Pratik Q. Deb, Luke Fritzky, Selvakumar Subbian, Nan Gao, Yungtai Lo, Mirella Salvatore, Amariliz Rivera, Alexander Lemenze, Patricia Fitzgerald-Bocarsly, Sanjay Tyagi, Wuyuan Lu, Aimee Beaulieu, Theresa L. Chang

## Abstract

Interferon ε (IFNε) is a unique type I IFN that has been implicated in host defense against sexually transmitted infections (STIs). Zika virus (ZIKV), an emerging pathogen, can infect the female reproductive tract (FRT) and cause devastating diseases, particularly in pregnant women. How IFNε contributes to protection against ZIKV infection *in vivo* is unknown. Here, we show that IFNε plays a critical role in host protection against vaginal ZIKV infection in mice. We found that IFNε was expressed not only by epithelial cells in the FRT, but also by certain immune and other cells at baseline or after exposure to viruses or specific TLR agonists. IFNε-deficient mice exhibited abnormalities in the epithelial border and underlying tissue in the cervicovaginal tract, and these defects were associated with increased susceptibility to vaginal, but not subcutaneous ZIKV infection. IFNε-deficiency resulted in an increase in magnitude, duration, and depth of ZIKV infection in the FRT. Critically, intravaginal administration of recombinant IFNε protected *Ifnε^-/-^* mice and highly susceptible *Ifnar1^-/-^*mice against vaginal ZIKV infection, indicating that IFNε was sufficient to provide protection even in the absence of signals from other type I IFNs and in an IFNAR1-independent manner. Our findings reveal a potentially critical role for IFNε in mediating protection against transmission of ZIKV in the context of sexual contact.

**Significance:** Interferon ε (IFNε), a unique Type I IFN that is highly expressed in the epithelium of the female reproductive tract (FRT), is thought to protect the host against sexually transmitted infections (STIs) but the mechanism of action is not defined. Zika virus (ZIKV), a causative agent for preterm birth and other severe diseases in pregnant women, can be spread through vaginal transmission. Here, we show that mice lacking the *Ifnε* gene have abnormal epithelial development and tissue architecture in the cervicovaginal tract. The role of IFNε in protecting host against ZIKV is FRT-specific and is independent of IFNAR1 signaling. Our findings suggest potential preventive strategies based on harnessing mucosal immunity against STIs.

## Introduction

Host defense against microbial invasion in the female reproductive tract (FRT) depends on maintenance of an effective epithelial barrier and continual surveillance by immune cells in the mucosa. Disruption of barrier function or immune responses in the FRT is associated with sexually transmitted infections (STIs) and other genitourinary infections (1-6). Zika virus (ZIKV) is an emerging flavivirus that is principally transmitted by Aedes mosquitoes, but can also be spread through vaginal transmission as an STI (7-11). ZIKV infection can cause severe adverse effects during pregnancy, ranging from preterm birth and miscarriage to microcephaly and other congenital defects in the fetus (12). ZIKV infection has also been associated with diverse pathologies in adults and children, including Guillain-Barre syndrome, myelitis, and neuropathy (13, 14). The cellular and molecular signals that mediate protection against ZIKV infection, particularly in the context of transmission through sexual contact, remain incompletely understood.

Prior studies in humans and mice have demonstrated an important role for Type I interferon (IFN) epsilon, IFNε, in protection against STIs including herpes simplex virus 2 (HSV2), ZIKV, and chlamydial infection (15-20) but the mechanism of anti-viral activities in vivo was not defined. Although IFNε is classified as a Type I IFN, it differs from other Type I IFNs, i.e., IFNα and IFNβ, in many respects, including its low (∼ 30%) amino acid homology and distinct expression patterns (15). Indeed, IFNε is constitutively expressed in epithelial cells lining many mucosal tissues including the FRT, whereas IFNα/β expression in these tissues is typically limited to settings of infection or inflammation (21). The mechanisms by which IFNε and other Type I IFNs mediate anti-viral defense also differ. We have previously shown that IFNε protects human primary macrophages against HIV infection via mechanisms distinct from IFNα2, through induction of reactive oxygen species (ROS), distinct pro-inflammatory cytokines and chemokines, and phagocytosis (22). Moreover, IFNε (at mid ng/ml ranges) has no or weak antiviral activities in transformed cell lines, while IFNα/β exhibit potent antiviral activities in diverse primary and transformed cell types (22-25).

In this study, we demonstrate a critical role for IFNε in host defense against vaginal ZIKV infection in mice. We found that IFNε-deficient mice are specifically susceptible to ZIKV when challenged via the intravaginal but not subcutaneous route, suggesting that IFNε plays a specific role in protecting against ZIKV transmission in the context of sexual contact. Our data indicate that IFNε is required to maintain normal epithelial and submucosal architecture in the cervicovaginal tissue of mice at steady state, suggesting that IFNε may promote resistance to ZIKV infection by maintaining tissue integrity. Notably, recombinant IFNε was sufficient to protect highly susceptible *Ifnar1^-/-^* mice from intravaginal ZIKV challenge, suggesting that IFNε provides host protection even in the absence of signals from other Type I IFNs, and it mediates this protection through IFNAR1-independent mechanisms. Taken together, our data highlight a novel role for IFNε in maintaining epithelial and tissue structure in the FRT and for mediating FRT-specific protection against ZIKV infection.

## Results

### IFNε is expressed by epithelial and non-epithelial cells in the FRT

To better understand steady-state expression patterns of IFNε *in vivo,* we first assessed *Ifnε* transcript abundance in different organs and mucosal tissues from naïve wild type (WT) mice. In agreement with a previous study (16), *Ifnε* expression was highest in FRT tissues compared to other tissues, although abundant *Ifnε* transcripts were also detected in the lung and intestinal mucosal tissues (**Figure 1A**). Within the FRT compartment, we found that *Ifnε* expression was highest in the cervicovaginal tissue (CVT), as compared to the uterus and ovaries. Consistent with our gene expression data, IFNε protein was readily detected in vaginal and uterine tissues from WT mice (**Figure 1B**).

**Figure 1.**
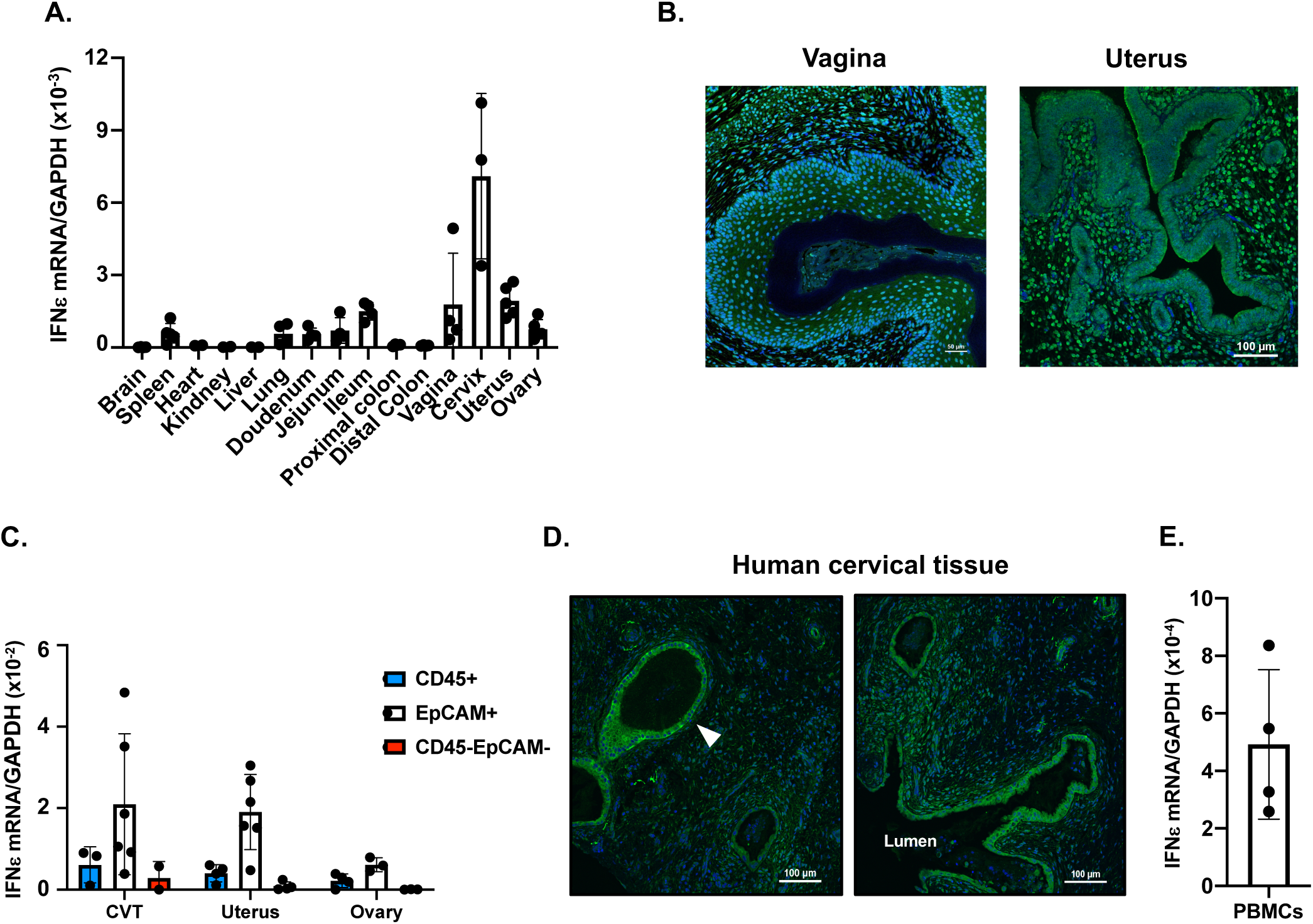
IFNε is expressed in epithelial and non-epithelial cells at the female reproductive tract. (A) Total RNAs were extracted from different tissues of wild-type (WT) mice. IFNε mRNA levels were quantified by RT-qPCR and normalized by GAPDH using the 2^-ΔΔ*CT*^ method. Each point represents one animal. (B) Expression of IFNε protein (Green) in vaginal (*left*) and uterus (*right*) tissue. Nuclei were stained with DAPI (blue). (C) IFNε mRNA levels were quantified by RTqPCR of total RNA from sorted CD45+ cells, epithelial cells (EpCAM), and CD45^-^EpCAM^-^ cells from cervicovaginal tissue (CVT), uterus, and ovary. (D) Detection of human IFNε proteins in human cervical tissues by immunofluorescence staining. The arrow indicates glandular epithelial cells expressing IFNε proteins. (E) Total RNAs were extracted from freshly isolated PBMCs from different donors. IFNε mRNA levels were quantified by RT-qPCR.

To identify specific cell types that express *Ifnε* transcripts in the FRT, we performed RT-qPCR on sort-purified CD45^+^ immune cells, EpCAM^+^ epithelial cells, and CD45^−^EpCAM^−^ cells from the ovaries, CVT, and uteri of WT mice. *Ifnε* transcripts were most abundant in epithelial cells, but were also detected in immune and non-immune CD45^−^EpCAM^−^ cells from the FRT (**Figure 1C**). These findings suggest that IFNε is expressed by diverse cell types in the FRT, including epithelial cells and immune cells.

Given our findings of IFNε expression by epithelial and immune cells in mouse FRT tissue, we assessed IFNε expression in primary human cervical tissue and human PBMCs. Consistent with our findings in mice, IFNε protein was readily detectable in the luminal and glandular epithelial cells and in non-epithelial cells distributed throughout the submucosa in cervical tissue from healthy donors (**Figure 1D**). We also observed low but detectable levels of *IFNε* transcripts in PBMCs from healthy donors (**Figure 1E**), suggesting that immune cells may also serve as a key source of IFNε in humans.

### IFNε is induced in response to TLR activation and viral infection

Prior studies reported that mouse bone marrow-derived macrophages do not express IFNε at baseline or in response to TLR activation or viral infection (16). However, whether inflammatory or infectious stimuli are sufficient to induce IFNε expression in other cell types remains an open question. To address this, we first investigated the impact of stimulation with TLR agonists on IFNε expression in freshly isolated human PBMCs. *IFNε* transcripts were low at baseline but were significantly upregulated in response to TLR3 or TLR4 activation (**Figure 2A**). Since activated plasmacytoid dendritic cells (pDCs) produce high levels of IFNα, we also measured *Ifnε* expression in human pDCs exposed to various viruses or cytokines for 4 h. *IFNε* transcripts were undetectable in pDCs at baseline, but were significantly upregulated in response to Sendai virus, HSV1, and Influenza virus (**Figure 2B**). In contrast, exposure to HIV did not induce detectable *IFNε* expression in pDCs. Cytokines were less effective than viruses at inducing *IFNε* expression in pDCs. IFNα induced very low levels of *IFNε* in pDCs from some but not all donors, whereas IFN-λ, IL-3, and IL-10 did not induce detectable *IFNε* expression in pDCs from any donor (**Figure 2B**).

**Figure 2.**
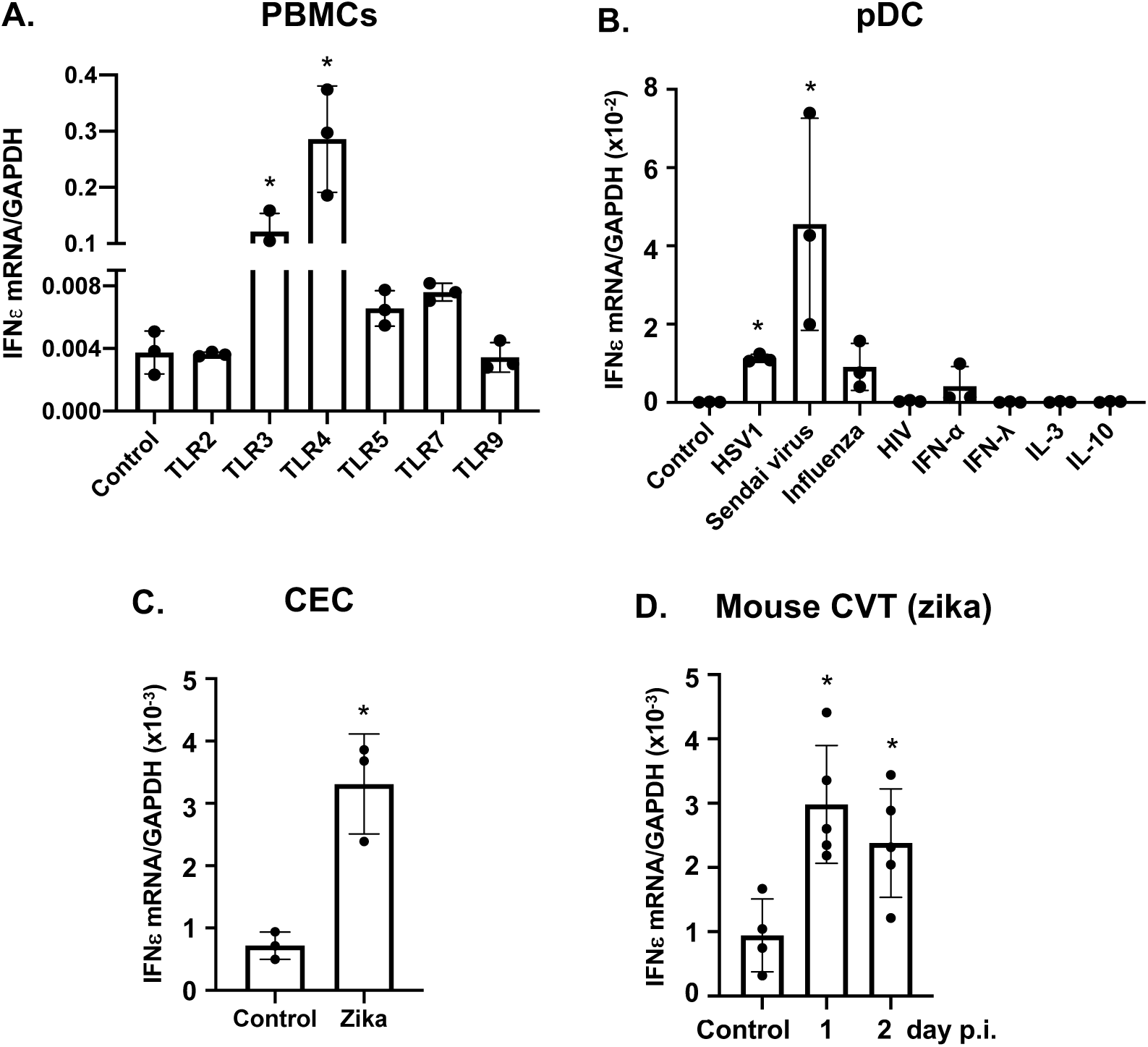
IFNε is induced in response to viral infection and TLR activation. (A) Freshly isolated PBMCs from different donors were stimulated with agonists for TLR2 (Pam3CSK4,100 ng/ml), TLR3 (poly (I:C), 1 μg/ml), TLR4 (LPS, 10 ng/ml), TLR5 (FLA-ST, 50 ng/ml), TLR7 (Imiquimod, 500 ng/ml), or TLR9 (ODN2006, 5 μM) for 6 h. Total RNAs were prepared and IFNε gene expression was quantified by RT-qPCR. (B) Plasmacytoid dendritic cells (pDCs) from different donors were purified using a negative selection pDC isolation kit. Purified pDCs were untreated (control) or treated with HSV1 2931, Sendai VR 907, influenza virus PR/8/34, HIV MN, rIFNα, IFNλ, IL-3, or IL-10 for 4 h. Total RNAs were prepared, and IFNε mRNA levels were quantified by RT-qPCR. (C) IFNε was determined by RT-qPCR in primary cervical epithelial cells (CEC) 24 h after ZIKV infection. (D) Induction of IFNε gene expression in the cervicovaginal tract of WT mice in response to ZIKV infection. \**p*<0.5, treated vs untreated controls.

Our studies (**Figure 1**) identified epithelial cells as key IFNε-expressing cell types in human and mouse FRT tissues at steady state. To determine whether viral infection further increased IFNε expression in epithelial cells, we measured *IFNε* transcript abundance in primary human cervical epithelial cells (CECs) cultured in medium or infected with ZIKV for 24 hours. Primary CECs expressed low but detectable levels of *IFNε* mRNA at baseline (**Figure 2C**), consistent with our findings of IFNε protein expression by epithelial cells in cervical tissue (**Figure 1D**). Nevertheless, infection with ZIKV elicited a marked increase in *IFNε* expression in CECs within 24 h post-infection (p.i.) (**Figure 2C**).

To determine whether IFNε expression in the FRT was increased by ZIKV infection *in vivo*, we challenged WT mice with ZIKV intravaginally and measured *Ifnε* transcript abundance in the CVT on day 1 and 2 p.i. These studies demonstrated a robust increase in *Ifnε* expression in the CVT in response to ZIKV infection (**Figure 2D**). Taken together, our data indicate that ZIKV and certain other viruses are strong inducers of IFNε expression in diverse cell and tissue types, including human PBMCs, pDCs, and CECs, and mouse CVT.

### IFNε-deficient mice exhibit abnormal epithelial borders and collagen deposition in the FRT

As a tool to investigate the role of IFNε in protective host responses *in vivo*, we generated *Ifnε^-/-^* mice using CRISPR/Cas9 technology (diagrammed in **Figure S1**). Successful disruption of the *Ifnε* gene was confirmed by genomic sequencing (data not shown), by loss of *Ifnε* transcripts (**Figure 3A**), and by loss of protein expression (**Figure 3B**). We used these mice to investigate the impact of IFNε-deficiency on global gene expression in the CVT to identify tissue-level processes regulated by IFNε signaling. Notably, genes identified as differentially expressed in CVT from naïve *Ifnε^-/-^* versus WT mice were significantly associated with pathways involved in tissue remodeling and/or epithelial barrier function, including tight junction signaling, fibrosis, epithelial adherens junction signaling, integrin signaling, and actin cytoskeleton signaling pathways (**Figure 3C**). Moreover, numerous genes encoding collagens, Wnt proteins, tight junction proteins, and chemokines or chemokine receptors were dysregulated in IFNε-deficient tissue (**Figure 3D** and Supplementary Information, Table S1).

**Figure 3.**
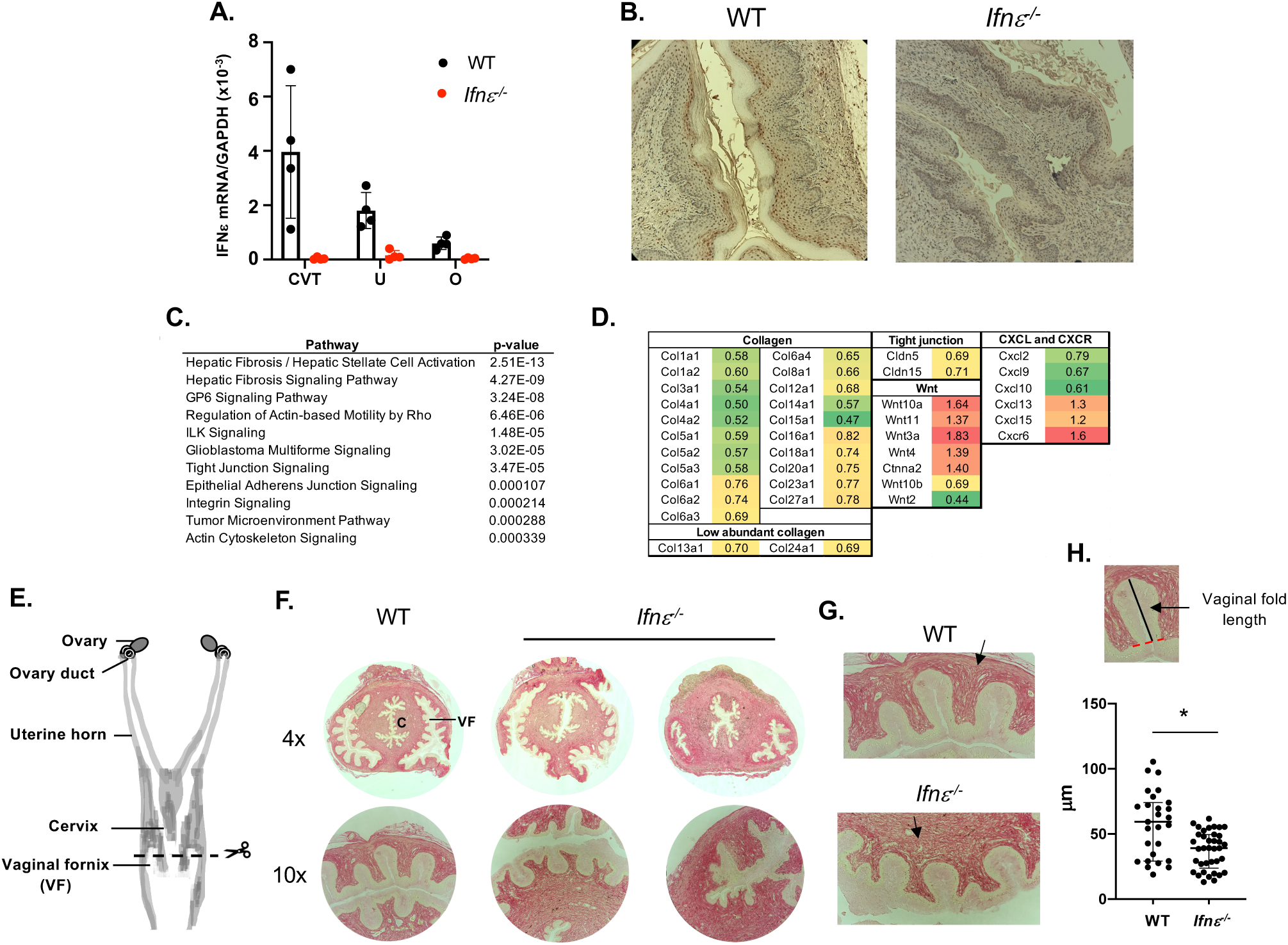
*Ifnε^-/-^* mice exhibit aberrant epithelial and tissue structure in the cervicovaginal tissue (CVT). (A) *Ifnε^-/-^* mice were generated using CRISPR/Cas9 technology. Knockdown of IFNε gene expression in CVT, uteri (U), and ovaries (O) of *Ifnε^-/-^* mice was confirmed by RT-qPCR. (B) IFNε protein expression in WT and *Ifnε^-/-^* mice was determined by IHC. IFNε proteins (brown) were found in the vaginal tissue of WT mice but not of *Ifnε^-/-^* mice. (C, D) WT and *Ifnε^-/-^* mice were synchronized to the diestrus stage by Depo-Provera. Total RNAs from the CVT were collected on day 12 after injection. RNAseq analyses showing pathways (C) and genes (D) involved in epithelial integrity, tissue structure, and wound healing in CVT of *Ifnε^-/-^* mice. (E) A schematic diagram of murine FRT. The site of sectioning for collagens and epithelial structure examination are indicated. (F) Collagens in the cervix of WT and *Ifnε^-/-^* mice were stained with Picro Sirius Red. Cervical canal (C) and vaginal fornix (VF) are indicated. (G) Higher power views (20x) of vaginal epithelial folds and surrounding collagen structure. Black arrows show the difference in collagen organization. (H) The length of vaginal epithelial folds in mid regions of vaginal epithelial fornix in WT and *Ifnε^-/-^* mice.

Wnt signaling critically regulates epithelial homeostasis and impacts the expression and function of collagens involved in tissue integrity and structure (26-31). Our finding of dysregulation of both collagen expression and Wnt signaling in CVT from *Ifnε^-/-^* mice raised the possibility that IFNε signaling may play a role in supporting proper tissue structure in the CVT. We therefore analyzed tissue architecture in the CVT of naive *Ifnε^-/-^* and WT mice (**Figures 3E-F**). These studies revealed discernible abnormalities in the vaginal fornix of *Ifnε^-/-^*mice compared to WT mice, including an underdeveloped epithelial border at the center of the vaginal fornix and disordered collagen deposition around the vaginal epithelial folds (10x) (**Figures 3F-G**). In addition, epithelial folds located in the central region of the vaginal fornix were significantly shorter in *Ifnε^-/-^* mice compared to WT mice (**Figures 3F-H**). These findings suggest that IFNε plays an important role in maintenance of optimal tissue architecture in the CVT, even in the absence of infection.

### IFNε is required for resistance to ZIKV infection following vaginal but not subcutaneous challenge

Our findings of abnormal barrier and tissue architecture in the CVT of *Ifnε^-/-^* mice led us to hypothesize that IFNε may specifically contribute to protection against pathogen entry in the CVT, a common transmission route for STIs. To test this, we challenged hormonally-synchronized *Ifnε^-/-^*, WT, and *Ifnar1^-/-^* mice with ZIKV PRVABC-59 via intravaginal or subcutaneous routes. Hormonal synchronization into diestrus was achieved by subcutaneous injection of Depo-Provera (Depo). In agreement with published data (9, 32-34), WT mice were more resistant to ZIKV infection than *Ifnar1^-/-^* mice regardless of the route of infection (**Figure 4A**). In contrast, *Ifnε^-/-^*mice were more susceptible to ZIKV infection than WT mice only when challenged via the vaginal route but not when challenged via the subcutaneous route (**Figure 4A**), supporting our hypothesis that IFNε provides specific protection against vaginal ZIKV transmission. Notably, increased viral loads in vaginally-infected *Ifnε^-/-^* mice were only observed in the CVT, whereas ZIKV mRNAs were similarly undetectable in the uteri and spleen of both *Ifnε^-/-^*and WT mice (**Figure 4B**). This finding contrasted with the high viral loads in all analyzed tissues from infected *Ifnar1^-/-^* mice (**Figure S2**). Viral RNAs were not detected in male *Ifnε^-/-^* mice or in non-Depo-treated female mice with subcutaneous ZIKV infection (**Figure S3**), indicating that the absence of detectable virus in subcutaneously-challenged *Ifnε^-/-^* mice was not an artifact of Depo treatment. Collectively, these findings highlight a specific role for IFNε in mediating protection against ZIKV when infection occurs via the vaginal route.

**Figure 4.**
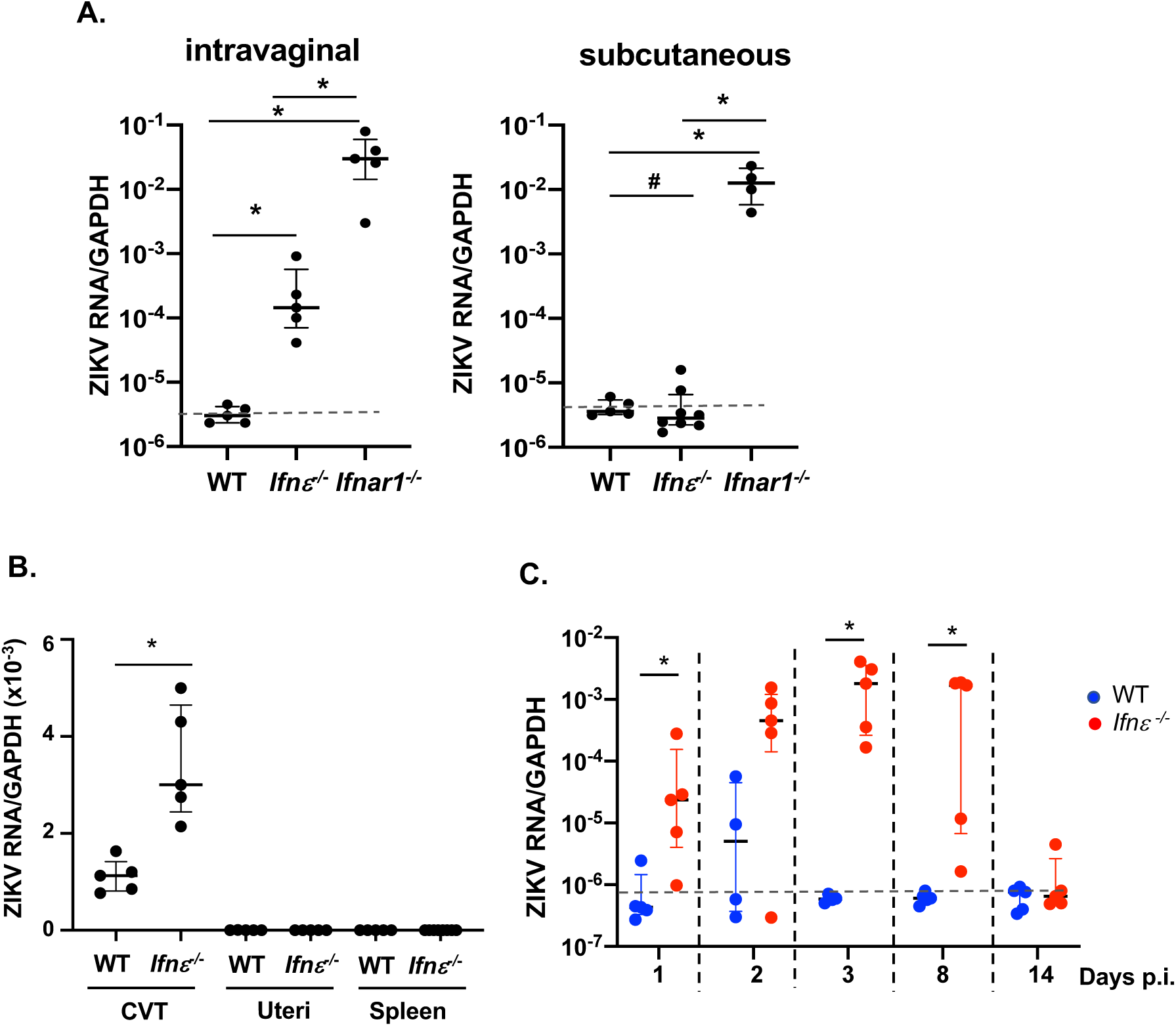
*Ifnε^-/-^* mice exhibit increased susceptibility to intravaginal ZIKV infection. (A) Depo-Provera-synchronized WT, *Ifnε^-/-^,* and *Ifnar1^-/-^* mice were challenged with ZIKV PRVABC59 through an intravaginal or subcutaneous route. Zika RNA levels at the CVT were determined by RT-qPCR at day 3 p.i. Data represent 5 independent experiments. (B) Depo-Provera-treated WT and *Ifnε^-/-^* mice were infected by zika virus intravaginally. Total RNAs from CVT, uteri, and spleen were harvested at day 3 p.i. Zika RNA levels were determined by RT-qPCR. Data represent 3 experiments (C) Depo-Provera-treated WT and *Ifnε^-/-^* mice were challenged with ZIKV intravaginally. Total RNAs of the CVT were prepared at different days p.i. ZIKV signals were determined by RT-qPCR. Data represent 3 experiments. The dash line indicates the background from uninfected mice. \**p<*0.05; ns, not significant.

To understand how IFNε signaling impacts the kinetics of ZIKV infection, we assessed viral loads in the CVT of vaginally-challenged WT and *Ifnε^-/-^*mice at different time points after infection. In WT mice, ZIKV RNAs were detectable in the CVT on day 1 and 2 p.i., but undetectable after day 3 p.i. (**Figure 4C**). In contrast, ZIKV RNA abundance was not only higher in the CVT of *Ifnε^-/-^* mice as early as day 1 p.i., but remained detectable until day 8 p.i. (**Figure 4C**).

We next used single molecule in situ hybridization (smFISH) to define the spatial parameters of ZIKV infection within the context of the architecture of the CVT on day 2 p.i. via the vaginal route. Although ZIKV RNA was readily evident in the mid-cervix in both WT and *Ifnε^-/-^* mice, it was only detected in the proximal cervix in *Ifnε^-/-^* mice but not WT mice (**Figures 5A-C**). In both the mid and proximal cervix of infected *Ifnε^-/-^* mice, ZIKV RNA was detected throughout the lamina propria and stroma. These findings suggest that IFNε may play a specific role in limiting the extent of viral spread in the FRT.

**Figure 5.**
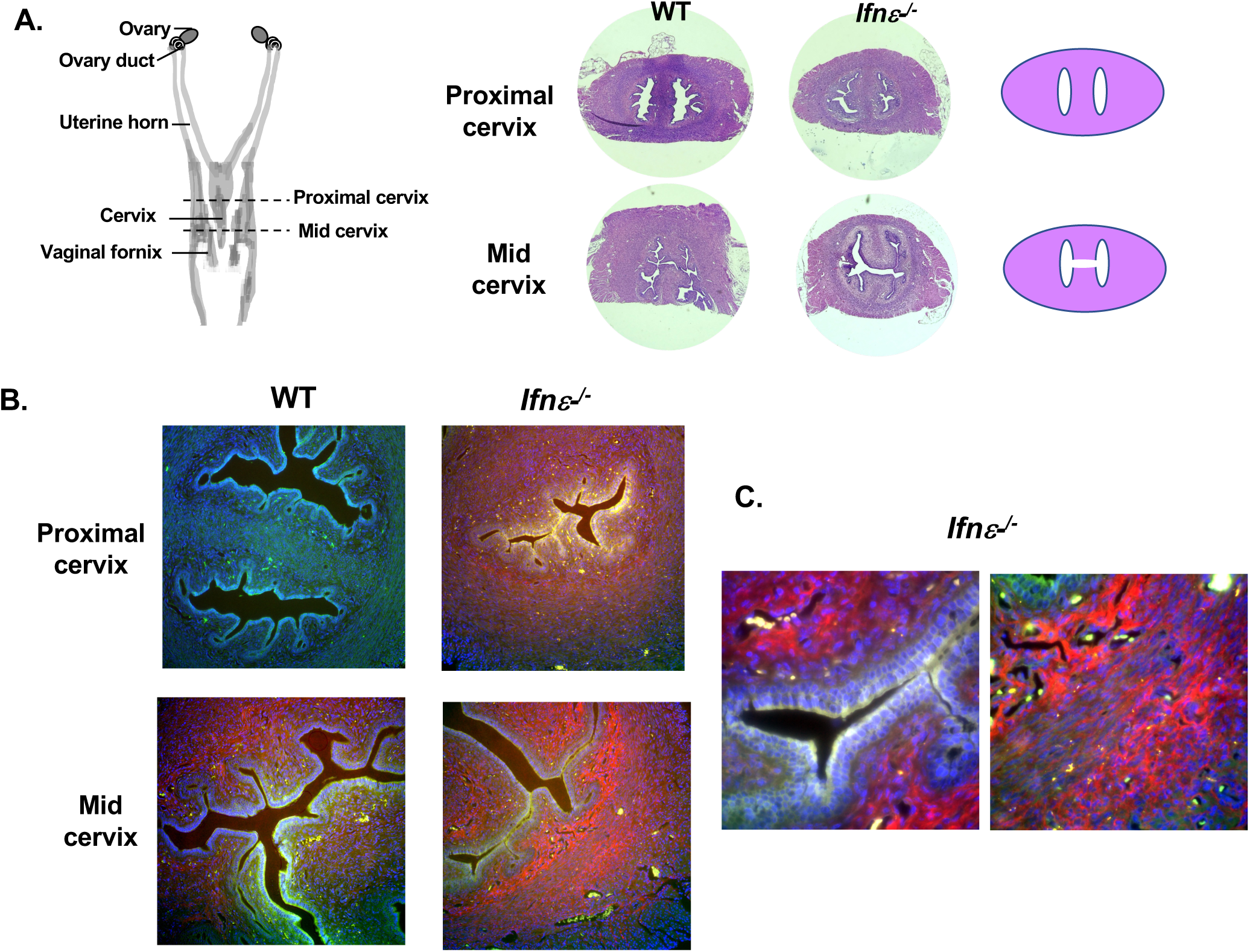
Faster dissemination of ZIKV of *Ifnε^-/-^* mice. (A) A serial sections of the CVT from uterus fundus toward the distal cervix. H&E staining of the proximal and mid cervix of WT and of *Ifnε^-/-^* mice used for smFISH are shown. (B). Adjacent sections to the H&E-stained sections were used for smFISH to detect ZIKV RNAs (red). Images at 20x are shown. Nuclei are in blue and autofluorescence is in green. Identical exposures were used for all images including both WT and *Ifnε^-/-^* mice. (C). Localization of ZIKA signals in *Ifnε^-/-^* mice (62x) in lamina propria (*left*) and stroma (*right*).

A role for IFNε in limiting the ascension of ZIKV toward the upper FRT was further supported by our observation that, while vaginal challenge caused mild genital erythema and inflammation of the CVT in both WT and *Ifnε^-/-^* mice (**Figures S4A-B**), tissue edema in the uterus was only observed in *Ifnε^-/-^* mice but not WT mice (**Figure S5**). Collectively, our findings highlight a specific function for IFNε in mediating protection against acute ZIKV infection via the vaginal route, and suggest that IFNε may act to restrict viral spread beyond the lower CVT.

To determine whether the increased susceptibility of *Ifnε^-/-^* mice to ZIKV was due to impaired antiviral responses, we assessed the expression of IFN-stimulated genes (ISGs) in the CVT from infected mice. ISGs including genes encoding IFNα and 2’5’ oligoadenylate synthetase (OAS) were upregulated to a comparable extent in the CVT of ZIKV-infected WT and *Ifnε^-/-^* mice (**Figure S6**). We also observed similar expression of *Ifnβ* transcripts in both cohorts, although total abundance at day 2 p.i was not significantly different from day 0. *Mx1* and *Isg15* were not induced. Taken together, these findings suggest that the increased susceptibility of IFNε mice to ZIKV is unlikely to be due to impaired ISG expression during acute infection.

### Recombinant mouse IFNρ, protein protects Ifnar1^-/-^ mice against ZIKV infection

Recombinant IFNε proteins have been shown to protect mice against intravaginal HSV2 and *chlamydia muridarum* infection (16). We therefore tested whether exogenous IFNε could protect mice against ZIKV infection. Vaginal administration of recombinant mouse IFNε protein protected *Ifnε^-/-^* mice against a subsequent intravaginal challenge with ZIKV (**Figure 6A**). Importantly, vaginal administration of recombinant IFNε, but not control linearized IFNε protein, was also sufficient to protect highly susceptible *Ifnar1^-/-^*mice from intravaginal ZIKV infection (**Figures 6B-C**), indicating that IFNε mediates protective effects through IFNAR1-independent mechanisms and is sufficient for protection even when signals from other Type 1 IFNs are absent. IFNε-mediated host protection was specific to vaginal infection because subcutaneous administration of recombinant IFNρ, did not protect *Ifnar1^-/-^* mice against subcutaneous ZIKV infection (**Figure 6C**). Additionally, IFNε proteins did not protect mice when administrated to ZIKV-infected mice on day 1 p.i. (**Figure 6D**). In summary, our data demonstrate that exogenous IFNε is sufficient to provide FRT-specific and IFNAR1-independent protection against ZIKV infection when administered prior to infection, even in highly susceptible *Ifnar1^-/-^* mice.

**Figure 6.**
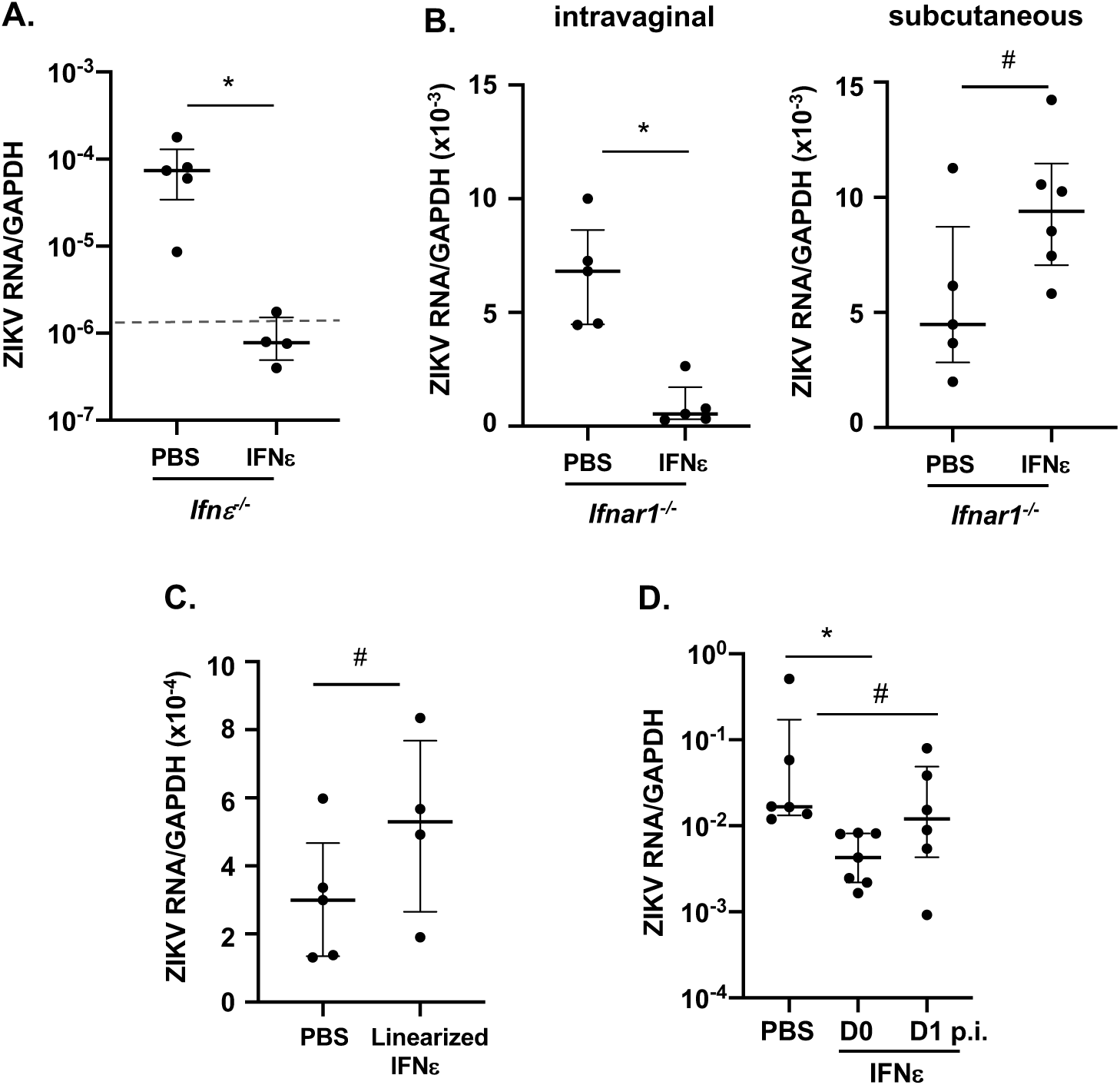
Intravaginal administration of murine IFN_ε_ proteins protect mice against ZIKV infection. (A) Depo-Provera synchronized *Ifnε^-/-^* mice were treated with recombinant mIFNε proteins (4 μg) through an intravaginal route for 6 h followed by ZIKV infection. ZIKV RNA levels at CVT were determined by RT-qPCR at day 3 p.i. The dash line indicates the background from uninfected control. (B) Depo-Provera synchronized *Ifnar1-/-* mice were treated with recombinant mIFNε proteins (4 μg) through an intravaginal or subcutaneous route for 6 h followed by zika infection via the same route. Zika RNA levels at the CVT and spleen were determined by RT-qPCR at day 3 p.i. (C) Depo-Provera-synchronized *Ifnar1-/-* mice were treated with PBS or linearized mIFNε proteins (4 μg) for 6 h intravaginally followed by intravaginal ZIKV infection. Viral RNA levels at the CVT were detected by RT-qPCR. (D) Depo-Provera-synchronized *Ifnar1-/-* mice were treated with mIFNε proteins (4 μg) intravaginally 6 h before or one day after (D1) intravaginal ZIKV infection. Viral RNA levels at the CVT were detected by RT-qPCR. *\*p<0.05, #p>0.05*. The values of the background from uninfected *Ifnar1-/-* mice were 1-3×10^-6^. Data represent 3-4 independent experiments.

## Discussion

IFNε was previously reported to protect mice against HSV2 and *Chlamydia muridarum* infection in the FRT but its mechanisms of action have remained incompletely understood (16). Our data suggest that IFNε may contribute to defense against microbial invasion in the FRT by ensuring maintenance of the epithelial barrier and tissue architecture in the CVT. Loss of IFNε disrupts this architecture and renders mice specifically susceptible to vaginal, but not subcutaneous, infection with ZIKV. In the context of vaginal infection, IFNε signaling not only limits the magnitude of ZIKV infection in the CVT, but also restricts viral persistence and dissemination beyond the lower FRT. Critically, exogenous IFNε is sufficient to protect not only *Ifnε^-/-^* mice but also highly susceptible *Ifnar1^-/-^*mice from intravaginal ZIKV infection. Thus, IFNε is sufficient to protect against vaginal ZIKV infection even when signals from other Type I interferons are absent. Taken together, our data suggest that IFNε serves as a critical gate keeper against ZIKV infection in the FRT, acting to prevent ascension of the virus into deeper tissue compartments that are critical for reproduction.

Our findings have direct relevance to ZIKV infections transmitted through sexual contact as opposed to transmission via mosquitoes. Within this context, *Ifnε^-/-^* mice represent a useful model for studying ZIKV transmission and pathogenesis in the CVT. Because of the CVT-specific role of IFNε in host defense, it will be possible to study early and local aspects of ZIKV pathogenesis that can be difficult to study in *Ifnar1*^-/-^ mice, which are highly immunodeficient and susceptible to uncontrolled viral replication and systemic dissemination (9, 32, 33).

A very recent report confirmed our findings of increased susceptibility of *Ifnε^-/-^* mice to intravaginal ZIKA infection (20). In this report, ZIKV was detected in vaginal wash but not vaginal tissues in ZIKV-infected *Ifnε^-/-^* mice at day 5 p.i. However, ZIKV was detected in uteri and ovaries (20). In our study, ZIKV was detected in the CVT but not tissues from upper FRT at day 3 p.i. The discrepancies may be due to differences in *Ifnε^-/-^* mouse lines, housing locations, duration of Depo-Provera treatment and ZIKV infection, and tissue preparation. A kinetic study on ZIKV signals of different regions of the FRT may provide insights into viral trafficking and clearance during the course of infection.

IFNε-mediated protection against viral infection *in vitro* is associated with induction of ISGs (21, 22). However, ISG expression in the CVT was similar in ZIKV-infected *Ifnε^-/-^*and WT mice, suggesting that ISG induction may not be required for the protective functions of IFNε in this setting, at least at early time points. Instead, our data suggest a novel function for IFNε in maintaining barrier integrity in the CVT. We found that IFNε-deficiency was associated with dysregulated expression of genes implicated in wound healing and epithelial barrier function in the CVT of naïve *Ifnε^-/-^*. These perturbations in gene expression were associated with abnormal collagen deposition and epithelial border structure in vaginal tissue nearest to the cervix. Taken together with our finding of exacerbated viral spread in the CVT of *Ifnε^-/-^* mice, these data suggest that IFNε may restrict viral dissemination by maintaining proper barrier function in the FRT.

The mechanisms by which IFNε modulates collagen and epithelial homeostasis in the FRT is unknown. Various cytokines, including Type I and II IFNs, TGF-β, and IL-1β, are known to modulate the synthesis of collagens required for maintenance of tissue integrity at steady state and during infection (35, 36). Other IFNs have been shown to regulate epithelial barrier function, including through effects on Wnt signaling (37-39). For example, IFNλ is constitutively expressed in gut and lung epithelial cells, where it regulates barrier integrity and mitigates inflammation-associated tissue damage during infection (40-43). We speculate that IFNε may exert similar functions in the epithelial and/or submucosal tissues of the FRT, and future studies on this topic are likely to deepen our understanding of the mechanisms by which IFNε protects against ZIKV and other sexually transmitted pathogens.

Prior studies reported high IFNε expression in epithelial cells and tissues in mouse and human FRT (16). Our data corroborate these findings, but further demonstrate IFNε expression by immune cells in the FRT of naïve mice, and by human peripheral blood immune cells activated with specific viruses or TLR ligands. In particular, Sendai virus strongly induced, and HSV1 moderately induced, IFNε expression in human pDCs from all donors, whereas influenza virus and HIV did not. Likewise, TLR 3 and 4 agonists, but not TLR 2, 5, 7, or 9 agonists, induced significant upregulation of IFNε expression in PBMCs. Our findings contrast with a prior work showing that TLR agonists do not induce IFNε in primary BMDMs, mouse embryonic fibroblasts, or a mouse macrophage cell line RAW264.7 (16). This same study reported that neither HSV2 nor *Chlamydial muridarum* infection drive increased IFNε expression in mouse uteri (16), whereas we found a significant increase in IFNε expression in CVT tissue in ZIKV-infected mice. The differences across studies suggest that IFNε expression is both cell type- and stimulus-specific. Moreover, the ability of certain viruses, but not others, to induce IFNε expression may depend on which pathogen-sensing pathway is engaged. In support of this, HSV1 induces IFNα expression through the TLR9-MyD88 pathway (44), while Sendai virus induces IFNα expression in pDCs via PKR-, TLR7/8-, and TLR9-independent pathways (45). The specific TLRs or pathogen-sensing pathways involved in Sendai- and HSV1-induced expression of IFNε in human pDCs or in ZIKV-induced mouse CVT remain to be determined.

Future studies are needed to determine whether IFNε produced by or acting on immune cells contributes to host protection in the FRT. However, IFNε has been shown to exert potent immunomodulatory effects on primary macrophages *in vitro* (22); to promote lymphocyte recruitment and protective CD8+ T cell responses in the lung during vaccinia virus infection (24); and to support NK cell recruitment in the FRT during chlamydia infection in mice (16). We speculate that IFNε signaling also regulates FRT immune responses to ZIKV infection. In support of this, our RNAseq data demonstrated altered chemokine and chemokine receptor expression in the CVT of naïve *Ifnε^-/-^* mice, highlighting a potential role for IFNε in regulating immune cell recruitment in the FRT even at steady state.

Although IFNε was previously reported to signal through IFNAR1 and IFNAR2 in BMDMs (16), we show here that exogenous mouse IFNε was sufficient to protect *Ifnar1^-/-^* mice against vaginal ZIKV infection, indicating that at least some anti-viral activities of IFNε in this setting are IFNAR1-independent. Moreover, these activities are structure-dependent as linearized IFNε protein failed to mediate protection.

In summary, we describe a novel role for IFNε in restricting viral replication and spread in the lower FRT during vaginal ZIKV infection in mice. We also demonstrate an FRT-specific role for IFNε in host defense against ZIKV infection, which is independent of IFNAR1 signaling, and we identify new cell and tissue type- and stimulus-specific features of IFNε expression in FRT and peripheral blood. Our findings offer insights into the potential use of IFNε to harness mucosal immunity for prevention of STIs.

## Methods

### Reagents

Histopaque®-1077, FBS, RPMI-1640, DMEM, and PBS were from Sigma-Aldrich (St. Louis, MO). Human IFNα2α was purchased from PBL Assay Science (Piscataway, NJ). Recombinant human IFN-λ1, IL-3, and IL-10 were purchased from PeproTech (Rocky Hill, NJ). Human IL-2 was from R&D Systems (Minneapolis, MN). Vero E6 cells were purchased from ATCC. Zika virus (PRVABC59 strain) was obtained through NIH Biodefense and Emerging Infections Research Resources Repository, NIAID, NIH. Zika virus was propagated in Vero E6 cells, and the virus titer was determined by plaque assays.

Murine IFNε proteins have three cysteine residues (C52, C162, C174), which can contribute to different disulfide linkages. Recombinant mIFNε with mutation of C174 to serine, a less conserved residue, had an improved capacity to induce ISGs (**Fig S7**) and was used in the studies. The DNA constructs used to express recombinant murine IFNε and IFNε analog with the C174S replacement in *E. coli* were synthesized and verified by Genescript (Piscataway, NJ). For bacterial expression, a codon-optimized IFNε cDNA was cloned into pET28. Inclusion bodies of IFNε expressed in *E. coli* BL21, prepared by Shenandoah Biotechnology, Inc. (Warminister, PA), were dissolved in 8 M GuHCl in the presence of DTT, purified by reversed-phase HPLC, lyophilized, folded, and verified by electrospray ionization mass spectrometry as described previously (22). Linear alkylated IFNε proteins were also prepared and verified by mass spectrometry as described previously (22). The endotoxin level in IFNε was below 0.01 ng/mL as determined by Pierce LAL Chromogenic Endotoxin Quantitation Kit (Thermo Fisher Scientific).

Rabbit polyclonal antibodies against human or murine IFNε proteins were generated by using peptides derived from IFNε protein sequences (Lampire Biological Laboratories (Pipersville, PA). The specificity of antibodies was determined by western blot analysis. Pre-immune sera were included as a control.

### Human cell isolation and cervical tissue collection

PBMCs were isolated from the blood of healthy human donors obtained from the New York Blood Center by Histopaque®-1077 gradient centrifugation. Plasmacytoid dendritic cells were isolated by negative selection from prepared PBMCs with the Human Plasmacytoid Dendritic Cell Isolation Kit-II from Miltenyi Biotec (Auburn, CA) according to the manufacturer’s instructions. The purity of enriched pDC (90-99%) was determined by flow cytometry (CD123 and BDCA-2 double positive). Stimulation of pDCs with viruses or cytokines were performed as described previously (46). IFNα2 (1000 IU/ml), IFN-λ1 (25 ng/ml), IL-3 (10 ng/ml), IL-10 (10 ng/ml), HSV-1 strain 2931 at a multiplicity of infection (MOI) of 1, influenza A virus PR/8/34 (4 hemagglutination U/ml), Sendai virus VR907 (16 hemagglutination U/ml), and (HIV-MN at 500 ng of p24/ml) were used to stimulate pDCs for 4 h.

Cervical tissues without gross pathology were obtained from women undergoing therapeutic hysterectomy. The study (Pro20140000108) was approved by Rutgers, New Jersey Medical School (NJMS) Institutional Review Board. Primary cervical epithelial cells were prepared as described previously (47).

### Mice

The protocols for animal handling were approved by the Institutional Animal Care and Use Committee at Rutgers, New Jersey Medical School (PROTO999900732). Animals were housed in individually ventilated and filtered cages under positive pressure in an SPF facility. C57BL/6J (WT) and *Ifnar1*^-/-^ mice on the same background were purchased from Jackson Laboratory (Bar Harbor, MA). *Ifnε^-^*^/-^ mice were generated on the C57BL/6J background by direct microinjection of CRISPR-Cas9 reagents into one-cell embryos. Founder 1 and 2 (**Figure S1**) were generated using 30 ng/μl Cas9 protein (PNABio), with 0.15 μM each of the crRNAs C110 CCTTGTACCACTCCAGTTCT and C112 ACTGAGAAGCAAGAGCCAAC along with 0.3 μM tracer RNA (Millipore-Sigma). Two founders were used to avoid confounding effects of disrupted gene sequences from CRISPR-Cas9. Both founders exhibited the same phenotypes. Founder 1 had a 373 bp deletion removing the sequence from 49 bp upstream of the initiating methionine to codon 109, eliminating 2/3 of the IFNε coding sequence. Translation of the recombinant resulted in a C-terminal 54 AA peptide. Founder 1 also had a 12 bp deletion in the 3’UTR. Founder 2 had a 588 bp deletion removing the entirety of the IFNε coding sequence from 48 bp upstream of the initiating methionine to 22 bp downstream of the stop codon. A 107 bp segment containing the coding sequence of the first 32 AA was inserted at the deletion site in an inverted orientation. No open reading frames containing any part of IFNε were present. Primers used to genotype Founders 1 and 2 were IFNEE 5’-CTGGAATGGGAACCAGAAAACCTAAG-3’ and IFNEF 5’-CTAGCCATCTTAGAACACAGTTAACC-3’. Founders were confirmed by Sanger sequencing of PCR fragments cloned into pCRTopo2.1 (Invitrogen). Transmitted alleles were confirmed in the F1 generation after crossing to C57BL/6J mice by Sanger sequencing.

To determine the effect of IFNε on host defense against Zika viruses, 8-12-week-old female mice were treated with 2 mg Depo-Provera (Depo) subcutaneously. Synchronization at the diestrus stage was confirmed on day 12-14 after Depo treatment. Mice were infected with 8×10^3^-1.5×10^4^ PFU of Zika virus through an intravaginal or subcutaneous route. Clinical scores were assessed based on the following criteria: 0, no apparent signs of disease; 1, genital erythema; 2, moderate genital inflammation and swelling; 3, mucus, swelling, and redness with presence of genital lesions; 4, severe genital lesions and/or hind limb paralysis as described by Fung et al (16). Fat tissues associated with the FRT were removed physically and the specific regions of FRT were harvested at indicated time points for RT-qPCR, smFISH, IHC, and collagen assays.

To determine IFNε expression in different cell types at the FRT, single-cell suspensions were generated as previously described (48). Briefly, the CVT, uteri, and ovaries were separated and minced with scissors, and then digested with 1 mg/mL collagenase A (Sigma-Aldrich) and 0.1 mg/mL DNase (Roche) in RPMI-1640 (Gibco) with shaking at 250 rpm at 37°C twice for 15 min. Tissues were mechanically dissociated by passage through an 18- or 20-gauge needle after each shaking cycle. Cells were filtered through 100 μm mesh and washed once with cold RPMI with 3% fetal bovine serum (FBS) (Fisher Scientific). Red blood cells were lysed with RBC lysis buffer (Tonbo Biosciences). Epithelial cells, CD45+ cells, and cells without either EpCam or CD45 markers were sorted using antibodies against EpCam (clone G8.8, Biolegend, San Diego, CA) or CD45 (clone 30-F11, eBiosciences) directly conjugated to a fluorophore for epithelial cells or hematopoietic cells, respectively, in a FACSAria II sorter (BD Biosciences). Cell purity was greater than 95%.

To prepare bone marrow-derived macrophages (BMDMs) from mice for testing murine IFNε activities, femurs were obtained from 8–12-week-old C57BL/6 mice. After euthanasia using CO_2_, the femurs were dissected using scissors, cutting through the tibia below the knee joints as well as through the pelvic bone close to the hip joint. After removing muscles connected to the bone, and the femurs were rinsed in ice-cold 70% ethanol for 1 min, washed with PBS, and then both epiphyses were removed. The bones were flushed with a 10mL syringe and a 27½ gauge needle with RPMI 1640 plus 10% FBS to extrude bone marrow, which was then homogenized by pipetting. Bone marrow cells were then collected by centrifugation and incubated in red blood cell lysis buffer at room temperature for 5 min. Cells (1×10^6^) were cultured in RPMI 1640 with 10% FBS, gentamicin, amphotericin, and murine M-CSF at 40 ng/ml in a 10 cm dish for 5-6 days to obtain BMDMs.

### Immunohistochemistry

Tissue sections of formalin-fixed, paraffin-embedded tissues were deparaffinized in xylene and rehydrated in a standard series of descending alcohol immersions. After antigen retrieval, sections were stained with antibodies (Abs) against IFNε (Rabbit anti-mouse IFNε, #24864) at a 1:400 dilution and incubated at room temperature for 50 min, according to the immunohistochemistry protocol provided in the Dako Envision Kit (Aligent). The images were taken using an ECHO microscope.

### Immunofluorescence microscopy

Tissue was mounted on slides, deparaffinized, and rehydrated. For antigen retrieval, tissue was brought to 90°C in 10 mM sodium citrate (pH 6.0) for 10 minutes. Slides were allowed to cool, then placed in PBS for 10 minutes. Tissue was blocked in 5% FBS for 30 minutes, then washed three times in PBS, 0.05% Triton for 5 min/wash. Primary antibody was diluted in SuperBlock (ThermoFisher) and applied to tissue overnight. Tissue was washed three times in PBS, 0.05%Trition for 5 min/wash. Secondary Alexa Fluor F(ab’)2 fragment antibody was diluted 1:1000 in Superblock and applied to tissue for 2 hours. Tissues were counterstained with DAPI and mounted using Vectashield mounting medium (Vector Labs, Burlingame, CA). Images were acquired on a Nikon Eclipse Ti confocal microscope and analyzed by NIS-Elements software.

### Collagen staining

Collagen assays were performed using tissues from WT and *Ifnρ.^-/-^* mice from the same experiment, which were stained at the same time to reduce variability. Tissue sections were deparaffinized using xylene, hydrated through graded ethanol, and washed with distilled Mili-Q water. Tissue slides were then incubated with Picro Sirius Red Stain Kit (ab150681, Abcam, Cambridge, UK) for 60 min at room temperature according to the manufacturer’s instructions. Slides were rinsed in Acetic Acid Solution twice, dehydrated with 100% ethanol, and then mounted using glycerol. Images were captured on an ECHO Revolve Microscope.

### Single molecule fluorescence in situ hybridization (smFISH)

Serial 4 μm sections of formalin-fixed, paraffin-embedded tissue were prepared by Nationwide Histology (Missoula, MT). smFISH for Zika virus RNA was performed following procedures described previously (49). A total of 48 3’-amino labeled oligonucleotide probes for the NS1 region of Zika virus strain PRVABC59 were designed using Stellaris Probe Designer with the highest stringency settings for human host and then synthesized by LGC Biosearch Technologies. Probes were pooled, labeled with Cy5, and purified by HPLC (50). The sequences of the probes are listed in Supplementary Table 2. Tissue slides were equilibrated in 2XSSC, 10% formamide in wash buffer and then hybridized in 50 µl hybridization buffer containing 25 ng of pooled probes overnight at 37°C in a humid chamber. The coverslips were washed twice for 5 min with wash buffer, equilibrated with 2XSSC supplemented with 0.4% glucose, and mounted using deoxygenated mounting medium supplemented with DAPI (49). Images were acquired using a Zeiss Axiovert M200 microscope.

### Real-time RT-qPCR

Mouse tissues were homogenized using soft tissue homogenizing beads (VWR, Radnor, PA). Total RNA was isolated using TRIzol® (Life Technologies, Carlsbad, CA). First-strand cDNA was synthesized by incubating 1000 ng total RNA with oligo(dT)_12-18_ (25 μg/ml) or random primers (2.5 μM), and dNTPs (0.5 mM) at 65°C for 5 min followed by quick-chilling on ice. Reverse transcription was performed at 42°C for 50 min and 70°C for 15 min using SuperScript III Reverse Transcriptase. The PCR reaction contained cDNA equivalent to 50 ng of RNA input, 200 nM primer sets, and SYBR Green Master Mix (QIAGEN, Valencia, CA), and was run on a StepOnePlus real-time PCR system (Life Technologies, Carlsbad, CA). PCR conditions were 95°C denaturation for 10 minutes, 40 cycles of 95°C for 15 seconds, 60°C for 60 seconds. PCR products were quantified and normalized relative to the amount of GAPDH cDNA amplified in the same tube. Relative quantification of gene expression was calculated using the 2^-ΔΔCt^ (Ct, threshold cycle of real-time PCR) method according to the formula: ΔCT=Ct_GAPDH_ –Ct_target_, ΔΔCt=ΔCt_control_ – ΔC_target_. Primer sequences are listed in Supplementary Table 3.

### RNA sequencing analysis

RNA sequencing from WT and *Ifnε^-/-^* mice were performed by Azenta using an Illumina sequencer. Analysis was performed in accordance with the nf-core RNA-sequencing guidelines (version 1.4.2). Briefly, the output reads were aligned to the GRCm38 genome using STAR (version 2.6.1d), followed by hit count generation using featureCounts (version 1.6.4) and StringTie (version 2.0). Read counts were normalized and compared for differential gene expression using DESeq2 (version 3.10) with significance at False Discovery Rate (FDR) adjusted p-value < 0.05. RNAseq data has been submitted to GEO (Accession number: GSE228359). To identify the pathways/networks and biological functions perturbed in *Ifnε^-/-^* mice compared to WT mice, the significant differentially expressed genes were analyzed using Ingenuity Pathway Analysis (IPA, INGENUITY, Qiagen, Redwood City, CA) as described (51). Fisher’s exact test was used to calculate p-values used to rank networks of SDEG; p < 0.05 was considered statistically significant.

### Statistical analysis

Statistical comparisons were performed using Mann Whitney U test or one-way ANOVA with Tukey’s post hoc tests; *p* < 0.05 was considered significant.

## Acknowledgements

We thank Amy Rosenfeld and Vincent Racaniello for the technical support on Zika viral infection assay, Pierre Lespinasse and Debra Heller for their support on acquiring human cervical tissues, Pam De Lacy and her team at Shenandoah Biotechnology, Inc. for production of IFNε proteins in *E. coli*, Peter Romanienko at Rutgers, genomic editing core facility for generating *Ifnε^-/-^* mice, Heather Marlatt at Nationwide histology for preparation of tissue sections, and Eric Milner for editing the manuscript. This work was supported by NIH grants R01AI36948 to T.L.C, and R01 CA227291 to S.T.

---

**Fig S1.**
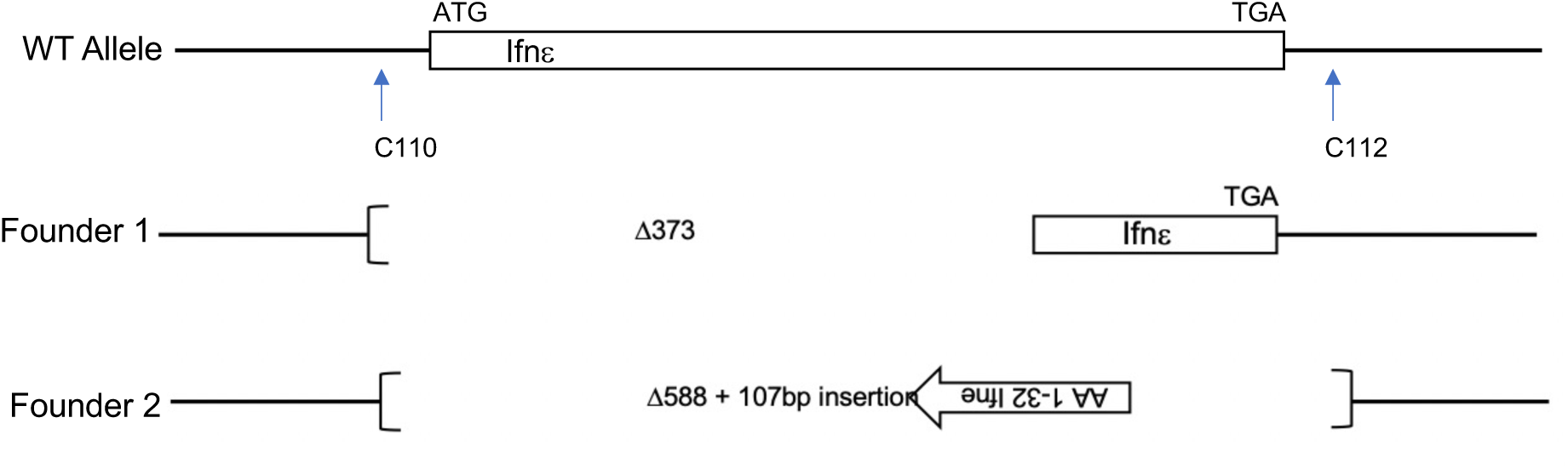
A schematic diagram of *Ifnε^-/-^* mice. *Ifnε^-/-^* mice were generated using CRISPR/Cas9 technology. The wild-type allele and the location of the guide RNAs (C110, C112) are shown. Founder 1 had a 373 bp deletion of the sequence from 49 bp upstream of the initiating methionine to AA 109, eliminating 2/3 of the IFNε coding sequence. Founder 1 also had a 12 bp deletion in the 3’UTR. Founder 2 had a 588 bp deletion of the entire IFNε coding sequence from 48 bp upstream of the initiating methionine to 22bp downstream of the stop codon. A 107 bp segment containing the coding sequence of the first 32 AA was inserted at the deletion site in an inverted orientation. No open reading frames containing any part of IFNε were present.

**Fig S2.**
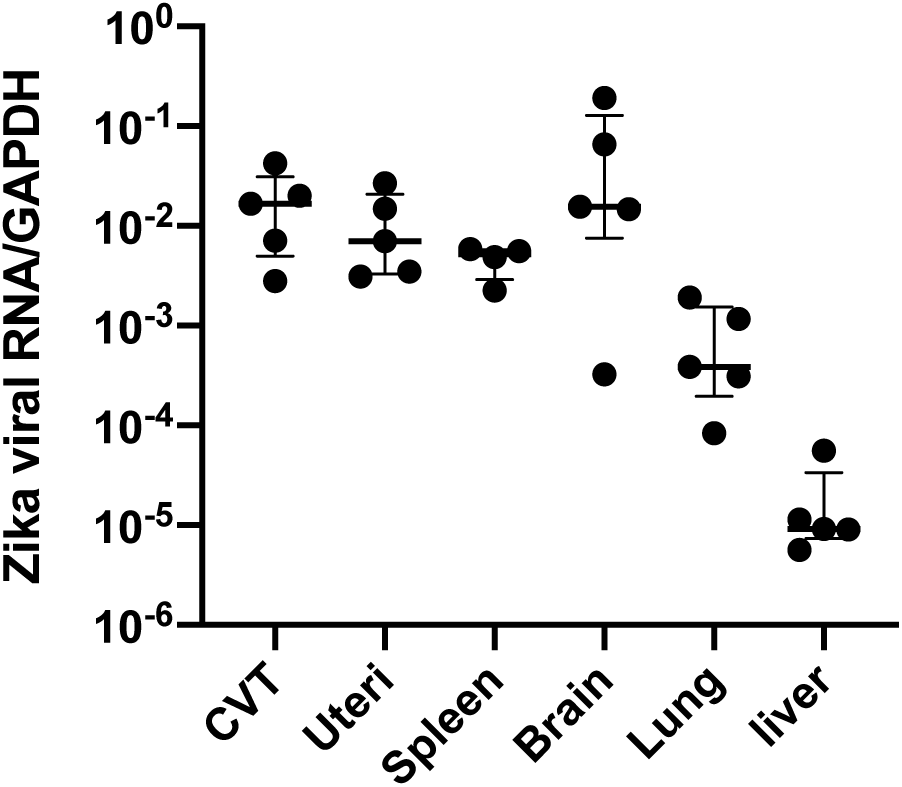
Zika viral RNAs were detectable in various tissues in *Ifnar1^-/-^* female mice. Depo-provera-treated *Ifnar1^-/-^* mice were infected by ZIKV via an intravaginal route. Various tissues were harvested at day 8 p.i. Total RNAs were prepared and Zika viral RNAs were analyzed by RT-qPCR.

**Fig S3.**
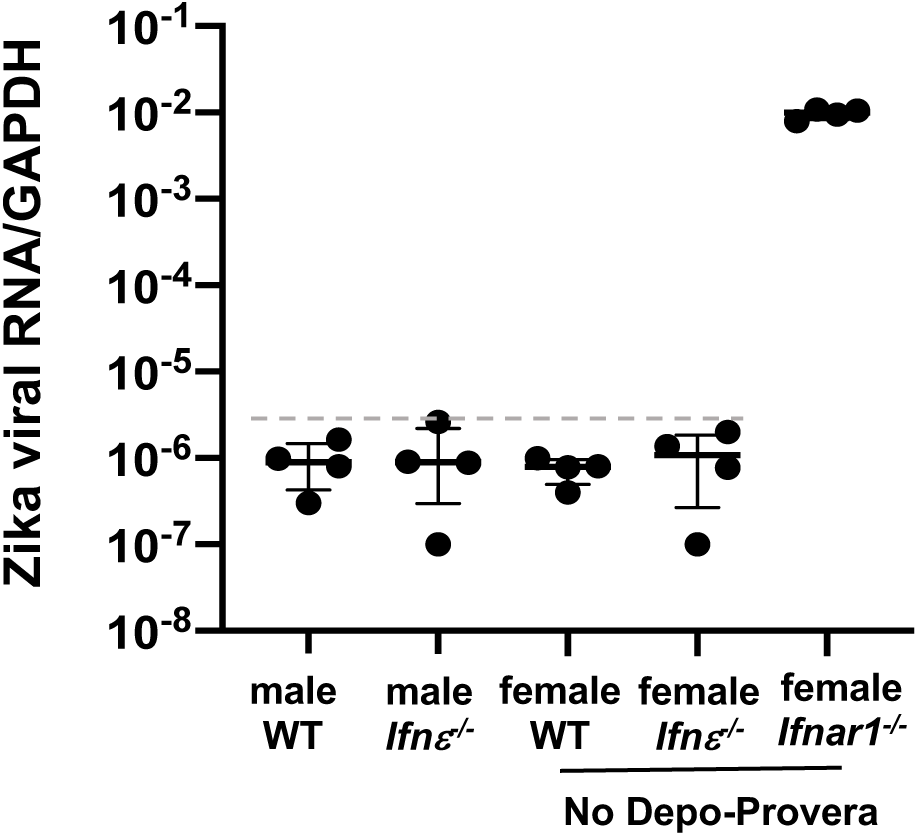
Zika viral RNAs were not detectable in the spleens of male or non-Depo-Provera treated female WT and *Ifnρ,^-/-^* mice subcutaneously infected with ZIKV. Male WT, male *Ifnρ,^-/-^*, female WT, female *Ifnρ,^-/-^*, and female *Ifnar1^-/-^* mice without Depo-Provera treatment were challenged with ZIKV subcutaneously. Spleens were harvested at day 8 p.i. ZIKV RNA levels were determined by RT-qPCR. (Dashed line indicates baseline signal from uninfected animals. Viral signals were not detectable in spleens of male or female WT and *Ifnρ,^-/-^* mice, whereas spleens from *Ifnar1^-/-^* mice harbored significant levels of ZIKV RNA.

**Fig S4.**
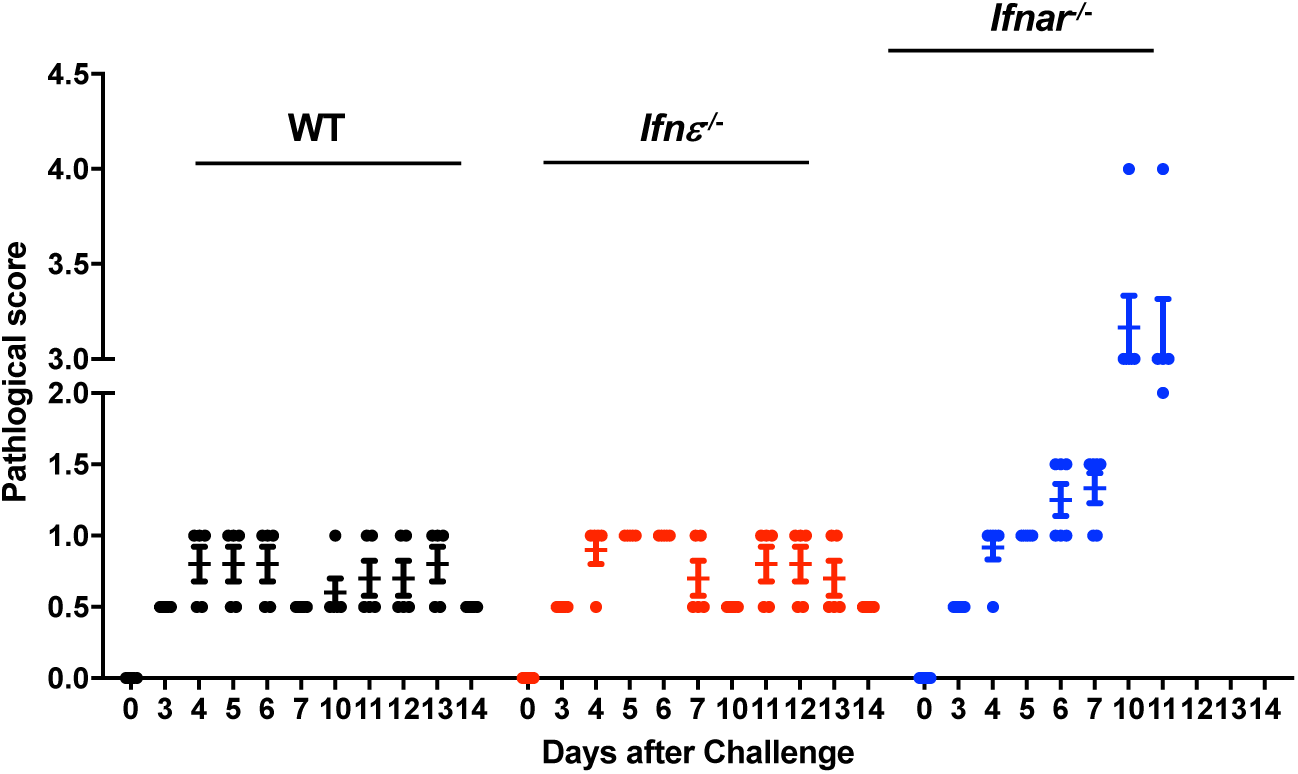
ZIKV-infected *Ifn*ε^−/*−*^ mice do not exhibit significant changes in clinical scores. Female WT, *Ifnρ,^-/-^*, and *Ifnar1^-/-^* mice were treated with Depo-Provera for 12-14 days before intravaginal ZIKV infection. *Ifnar1^-/-^* mice were included as a comparison. Clinical scores were recorded. The criteria of clinal score were listed in the method part. Higher score indicates more severe condition in mice. There were no significant differences in clinical scores or body weight loss between WT and *Ifnρ,^-/-^* mice. *Ifnar1^-/-^* mice exhibited significant pathology.

**Fig S5.**
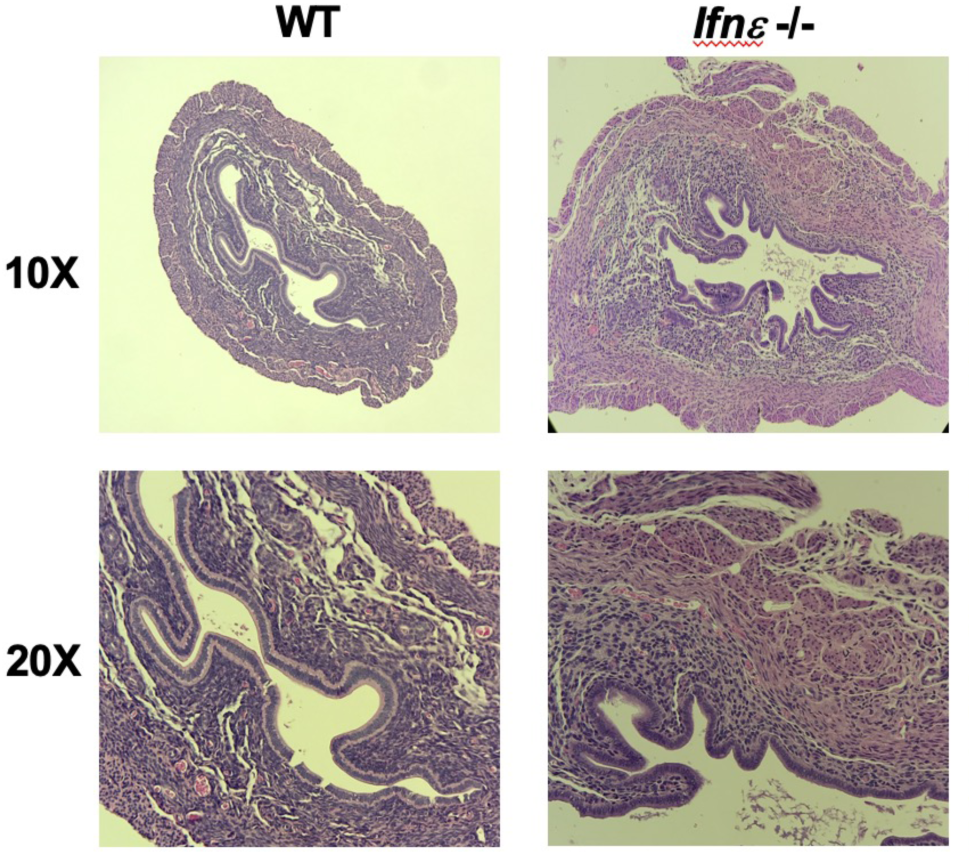
ZIKV induce tissue edema in the uterus of *Ifnε^-/-^*mice. Depo-Provera-synchronized WT and *Ifnε^-/-^* mice were infected with ZIKV. The FRT was harvested on day 8 p.i. and were examined by H&E staining.

**Fig S6.**
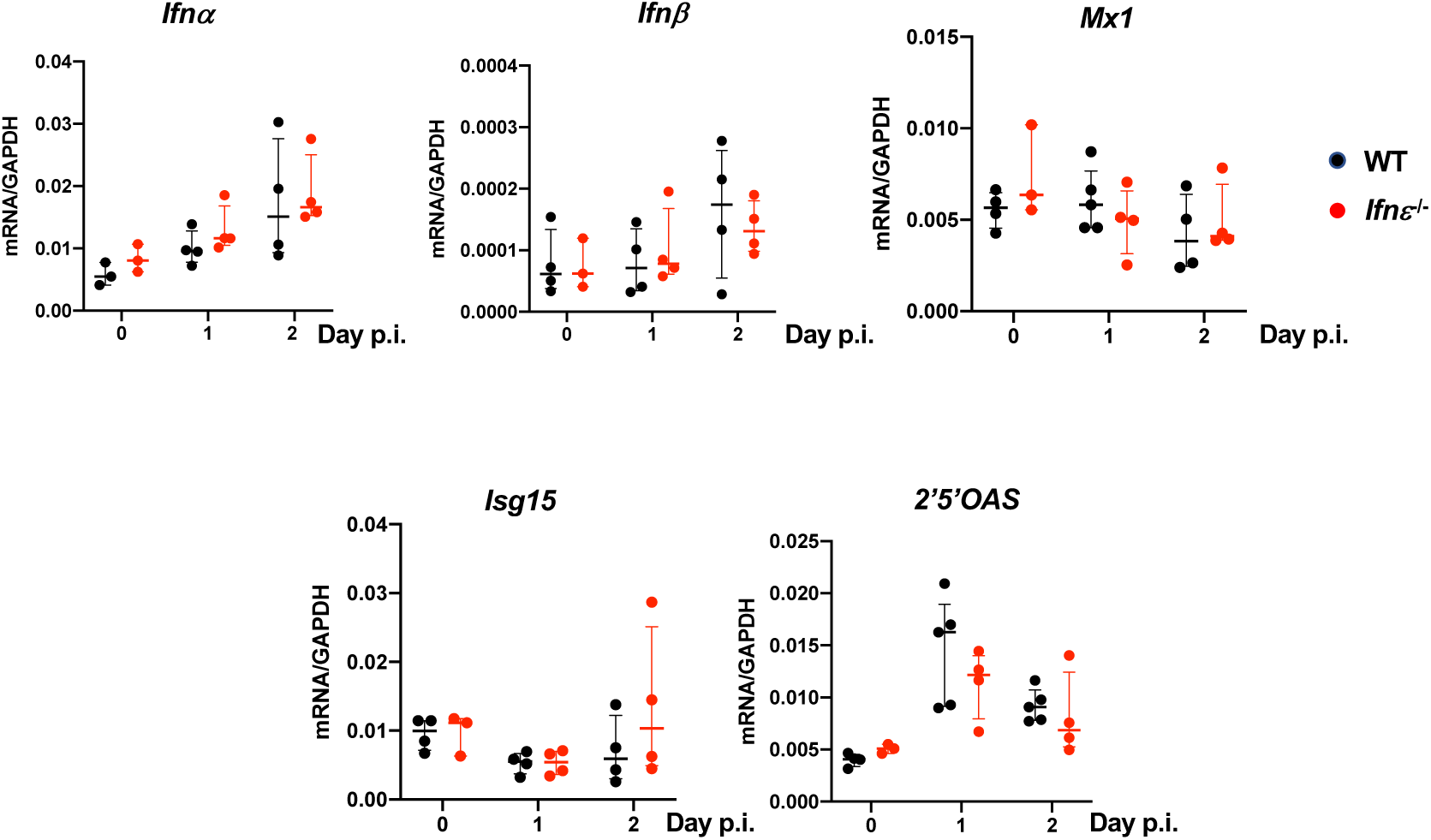
Induction of ISGs in the CVT of WT and *Ifn*ε^−/*−*^ mice in response to intravaginal ZIKV infection. WT and *Ifn*ε^−/*−*^ mice were infected with ZIKV intravaginally. Total RNAs from the CVT were prepared, and expression of ISGs was determined by RT-qPCR. *p<0.05 between WT and *Ifn*ε^−/*−*^ mice.

**Fig S7.**
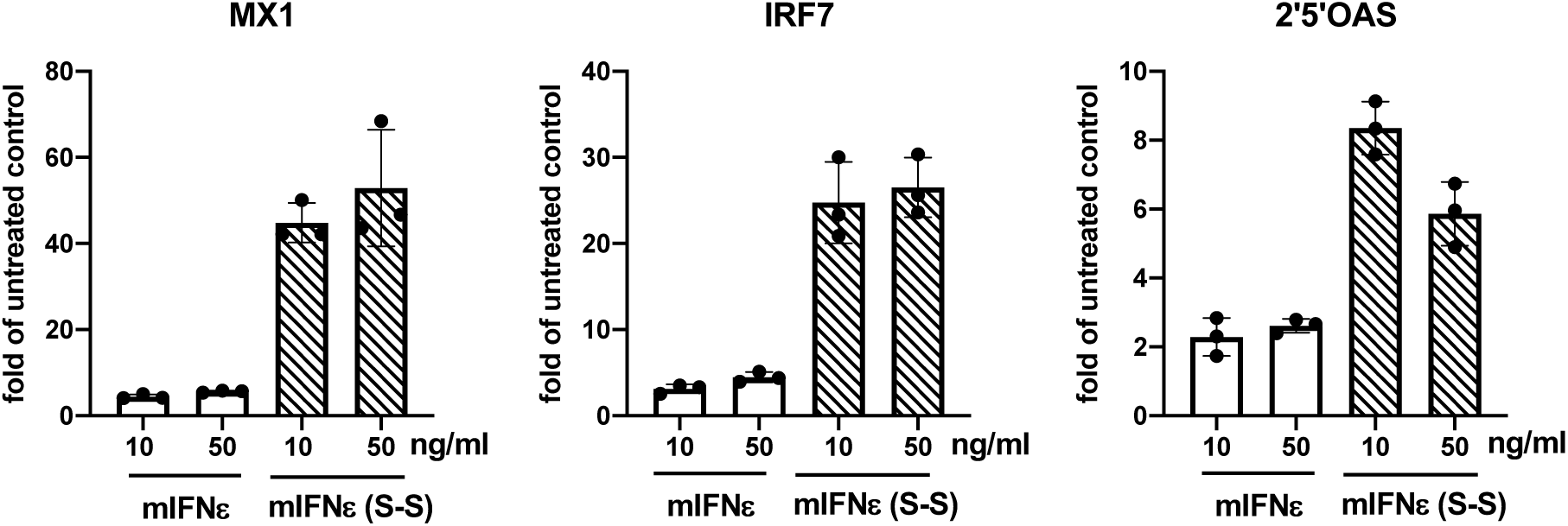
Recombinant mIFNε (S-S) proteins induce higher levels of ISGs. Recombinant mIFNε proteins and their analog, mIFNε (S-S) with a replacement of Cys 174 with serine to stabilize the protein structure. Bone marrow derived macrophages were treated with mIFNε proteins or mIFNε (S-S) at 10 or 50 ng/ml for 6 h. Total RNAs were prepared and the expression of ISGs including MX1, IRF7, and 2’5’OAS were analyzed by RT-qPCR.

