## Supplemental Table 1-3 for "Interferon ε restricts Zika virus infection in the female reproductive tract"

### Supplementary Table 1. Gene lists for Top pathways regulated by IFNε

| Ingenuity Canonical Pathways | -log(p-value) | Molecules |
|---|---|---|
| Hepatic Fibrosis /Hepatic Stellate Cell Activation | 12.6 | COL15A1,COL1A1,COL1A2,COL3A1,COL4A1,COL4A2,COL5A1,COL5A2,COL5A3,EDNRB,LEP,MYH10,MYH11,MYH14,MYL9,PDGFD PDGFRA,PDGFRB |
| Hepatic Fibrosis Signaling Pathway | 8.37 | COL1A1,COL1A2,COL3A1,COL5A3,FOS,GLI1,ITGB3,LEP,MYL9,MYLK,PDGFD,PDGFRA,PDGFRB,RHOJ,RHOV,RND3,WNT10A, WNT2,WNT3A |
| GP6 Signaling Pathway | 7.49 | COL15A1,COL1A1,COL1A2,COL3A1,COL4A1,COL4A2,COL5A1,COL5A2,COL5A3,ITGB3,LAMA2 |
| Apelin Liver Signaling Pathway | 6.87 | APLNR,COL1A1,COL1A2,COL3A1,COL5A3,PDGFRB |
| Regulation of Actin-based Motility by Rho | 5.19 | ACTC1,ITGB3,MYL9,MYLK,Ppp1r12b,RHOJ,RHOV,RND3 |
| ILK Signaling | 4.83 | ACTC1,FOS,ITGB3,MYH10,MYH11,MYH14,MYL9,RHOJ,RHOV,RND3 |
| Glioblastoma Multiforme Signaling | 4.52 | PDGFD,PDGFRA,PDGFRB,RHOJ,RHOV,RND3,WNT10A,WNT2,WNT3A |
| Tight Junction Signaling | 4.46 | ACTC1,FOS,MYH10,MYH11,MYH14,MYL9,MYLK,NECTIN1,SNAP25 |
| Cellular Effects of Sildenafil (Viagra) | 4.43 | ACTC1,MYH10,MYH11,MYH14,MYL9,MYLK,PDE5A,Ppp1r12b |
| Epithelial Adherens Junction Signaling | 3.97 | ACTC1,JUP,MYH10,MYH11,MYH14,MYL9,NECTIN1,SORBS1 |
| Integrin Signaling | 3.67 | ACTC1,ITGA11,ITGB3,MYL9,MYLK,Ppp1r12b,RHOJ,RHOV,RND3 |
| Tumor Microenvironment Pathway | 3.53 | COL1A1,COL1A2,FOS,HLA-A,ITGB3,LEP,PDGFD,SLC2A1 |
| Actin Cytoskeleton Signaling | 3.47 | ACTC1,ITGB3,MYH10,MYH11,MYH14,MYL9,MYLK,PDGFD,Ppp1r12b |
| PAK Signaling | 3.27 | ITGB3,MYL9,MYLK,PDGFD,PDGFRA,PDGFRB |
| Intrinsic Prothrombin Activation Pathway | 3.17 | COL1A1,COL1A2,COL3A1,COL5A3 |
| Signaling by Rho Family GTPases | 3.13 | ACTC1,CDH17,FOS,ITGB3,MYL9,MYLK,RHOJ,RHOV,RND3 |
| Sphingosine-1-phosphate Signaling | 3.05 | PDGFD,PDGFRA,PDGFRB,RHOJ,RHOV,RND3 |
| Osteoarthritis Pathway | 2.9 | ANXA2,DDIT4,GLI1,ITGB3,LEP,PTH1R,RARRES2,WNT3A |
| Atherosclerosis Signaling | 2.86 | COL1A1,COL1A2,COL3A1,COL5A3,F3,PDGFD |
| Human Embryonic Stem Cell Pluripotency | 2.73 | PDGFD,PDGFRA,PDGFRB,WNT10A,WNT2,WNT3A |
| RhoGDI Signaling | 2.64 | ACTC1,CDH17,ITGB3,MYL9,RHOJ,RHOV,RND3 |
| Colorectal Cancer Metastasis Signaling | 2.52 | FOS,PTGER3,RHOJ,RHOV,RND3,WNT10A,WNT2,WNT3A |
| Synaptogenesis Signaling Pathway | 2.51 | CDH17,EFNA3,MAP1B,NECTIN1,SNAP25,SYN2,SYT4,SYT5,THBS2 |
| Calcium Signaling | 2.44 | ACTC1,CASQ2,HTR3A,MYH10,MYH11,MYH14,MYL9 |
| Thrombin Signaling | 2.41 | GATA6,MYL9,MYLK,Ppp1r12b,RHOJ,RHOV,RND3 |
| Basal Cell Carcinoma Signaling | 2.31 | GLI1,WNT10A,WNT2,WNT3A |
| Caveolar-mediated Endocytosis Signaling | 2.28 | ACTC1,HLA-A,ITGA11,ITGB3 |
| Glioma Invasiveness Signaling | 2.28 | ITGB3,RHOJ,RHOV,RND3 |
| CXCR4 Signaling | 2.27 | EGR1,FOS,MYL9,RHOJ,RHOV,RND3 |
| Germ Cell-Sertoli Cell Junction Signaling | 2.22 | ACTC1,JUP,RHOJ,RHOV,RND3,SORBS1 |
| Tec Kinase Signaling | 2.2 | ACTC1,FOS,ITGB3,RHOJ,RHOV,RND3 |
| RhoA Signaling | 2.19 | ACTC1,MYL9,MYLK,Ppp1r12b,RND3 |
| Cdc42 Signaling | 2.16 | FOS,HLA-A,ITGB3,MYL9,MYLK,Ppp1r12b |
| Actin Nucleation by ARP-WASP Complex | 2.13 | ITGB3,RHOJ,RHOV,RND3 |
| Dendritic Cell Maturation | 2.07 | COL1A1,COL1A2,COL3A1,COL5A3,HLA-A,LEP |
| Regulation Of The Epithelial Mesenchymal Trans | 2.07 | GLI1,WNT10A,WNT2,WNT3A |

| | | |
|---|---|---|
| PDGF Signaling | 2.04 | FOS,PDGFD,PDGFRA,PDGFRB |
| Acetyl-CoA Biosynthesis III (from Citrate) | 2.02 | ACLY |
| Extrinsic Prothrombin Activation Pathway | 2 | F3,TFPI |
| Regulation of the Epithelial-Mesenchymal Transit | 1.99 | EGR1,PDGFD,PDGFRB,WNT10A,WNT2,WNT3A |
| Agranulocyte Adhesion and Diapedesis | 1.97 | ACTC1,CD34,MYH10,MYH11,MYH14,MYL9 |
| IL-8 Signaling | 1.91 | FOS,ITGB3,MYL9,RHOJ,RHOV,RND3 |
| Molecular Mechanisms of Cancer | 1.83 | FOS,GLI1,ITGB3,RHOJ,RHOV,RND3,WNT10A,WNT2,WNT3A |
| Sumoylation Pathway | 1.77 | FOS,RHOJ,RHOV,RND3 |
| IGF-1 Signaling | 1.76 | FOS,GRB10,IGFBP7,SFN |
| PPAR Signaling | 1.74 | FOS,PDGFD,PDGFRA,PDGFRB |
| Role of Osteoblasts, Osteoclasts and Chondrocyt | 1.74 | COL1A1,FOS,ITGB3,WNT10A,WNT2,WNT3A |
| HOTAIR Regulatory Pathway | 1.73 | COL1A1,COL1A2,WNT10A,WNT2,WNT3A |
| Semaphorin Signaling in Neurons | 1.71 | RHOJ,RHOV,RND3 |
| PCP pathway | 1.71 | WNT10A,WNT2,WNT3A |
| HMGB1 Signaling | 1.68 | FOS,LEP,RHOJ,RHOV,RND3 |
| cAMP-mediated signaling | 1.65 | APLNR,NPR3,PDE5A,PTGER3,PTH1R,RGS2 |
| Wnt/β-catenin Signaling | 1.6 | SOX21,SOX7,WNT10A,WNT2,WNT3A |
| Cholecystokinin/Gastrin-mediated Signaling | 1.57 | FOS,RHOJ,RHOV,RND3 |
| Role of NANOG in Mammalian Embryonic Stem | 1.57 | GATA6,WNT10A,WNT2,WNT3A |
| D-glucuronate Degradation I | 1.55 | CRYL1 |
| Acute Phase Response Signaling | 1.54 | CP,CRABP2,FOS,RBP2,SAA1 |
| Regulation Of The Epithelial Mesenchymal Trans | 1.47 | EGR1,FOS,PDGFD,PDGFRA,PDGFRB |
| Production of Nitric Oxide and Reactive Oxygen S | 1.46 | FOS,PPP1R14C,RHOJ,RHOV,RND3 |
| Retinoate Biosynthesis II | 1.43 | RBP2 |
| Sertoli Cell-Sertoli Cell Junction Signaling | 1.42 | ACTC1,ITGB3,JUP,NECTIN1,SORBS1 |
| Role of Wnt/GSK-3β Signaling in the Pathogenesi | 1.42 | WNT10A,WNT2,WNT3A |
| Phagosome Formation | 1.42 | ITGB3,RHOJ,RHOV,RND3 |
| Natural Killer Cell Signaling | 1.39 | COL1A1,COL1A2,COL3A1,COL5A3,HLA-A |
| Retinoate Biosynthesis I | 1.38 | ALDH1A2,RBP2 |
| Phospholipase C Signaling | 1.37 | ITGB3,MYL9,Ppp1r12b,RHOJ,RHOV,RND3 |
| Coagulation System | 1.36 | F3,TFPI |
| Semaphorin Neuronal Repulsive Signaling Pathw | 1.36 | ITGB3,MYL9,Ppp1r12b,VCAN |
| Citrulline-Nitric Oxide Cycle | 1.33 | ASS1 |
| Axonal Guidance Signaling | 1.32 | ADAMTS9,EFNA3,GLI1,ITGB3,MYL9,PDGFD,WNT10A,WNT2,WNT3A |
| HIPPO signaling | 1.32 | FAT4,PPP1R14C,SFN |
| G-Protein Coupled Receptor Signaling | 1.32 | APLNR,NPR3,PDE5A,PTGER3,PTH1R,RGS2 |
| Arginine Biosynthesis IV | 1.25 | ASS1 |
| Urea Cycle | 1.25 | ASS1 |
| Gαq Signaling | 1.2 | RGS2,RHOJ,RHOV,RND3 |
| Apelin Cardiomyocyte Signaling Pathway | 1.16 | APLNR,MYL9,MYLK |
| Virus Entry via Endocytic Pathways | 1.13 | ACTC1,HLA-A,ITGB3 |
| Leucine Degradation I | 1.08 | BCAT1 |

| | | |
|---|---|---|
| Paxillin Signaling | 1.08 | ACTC1,ITGA11,ITGB3 |
| Glioma Signaling | 1.06 | PDGFD,PDGFRA,PDGFRB |
| Protein Kinase A Signaling | 1.05 | MYH10,MYL9,MYLK,PDE5A,PPP1R14C,PTPRH,SFN |
| Prostanoid Biosynthesis | 1.04 | HPGDS |
| Ketolysis | 1.04 | BDH1 |
| Role of Tissue Factor in Cancer | 1 | EGR1,F3,ITGB3 |
| Ketogenesis | 1 | BDH1 |
| Systemic Lupus Erythematosus In T Cell Signalin | 1 | FOS,HLA-A,LEP,RHOJ,RHOV,RND3 |
| Role of PKR in Interferon Induction and Antiviral R | 0.987 | FOS,PDGFD,PDGFRB |
| Ephrin Receptor Signaling | 0.971 | EFNA3,ITGB3,PDGFD,SORBS1 |
| IL-15 Production | 0.963 | PDGFRA,PDGFRB,PEAK1 |
| Maturity Onset Diabetes of Young (MODY) Signa | 0.955 | ABCC9,FOXA2 |
| Clathrin-mediated Endocytosis Signaling | 0.947 | ACTC1,HIP1,ITGB3,PDGFD |
| RAR Activation | 0.943 | ALDH1A2,CRABP2,FOS,RBP2 |
| CREB Signaling in Neurons | 0.932 | ADGRA2,ADGRD1,APLNR,EDNRB,GPR153,PDGFRA,PDGFRB,PTGER3,PTH1R |
| Gαi Signaling | 0.932 | APLNR,NPR3,PTGER3 |
| Eicosanoid Signaling | 0.886 | HPGDS,PTGER3 |
| Superpathway of Citrulline Metabolism | 0.873 | ASS1 |
| PTEN Signaling | 0.854 | ITGB3,PDGFRA,PDGFRB |
| mTOR Signaling | 0.851 | DDIT4,RHOJ,RHOV,RND3 |
| Iron homeostasis signaling pathway | 0.848 | CP,PDGFRA,PDGFRB |
| Agrin Interactions at Neuromuscular Junction | 0.845 | ACTC1,LAMA2 |
| Ovarian Cancer Signaling | 0.833 | WNT10A,WNT2,WNT3A |
| Insulin Receptor Signaling | 0.833 | ACLY,GRB10,PPP1R14C |
| ERK5 Signaling | 0.824 | FOS,SFN |
| Isoleucine Degradation I | 0.824 | BCAT1 |
| Histamine Degradation | 0.824 | ALDH1A2 |
| Aryl Hydrocarbon Receptor Signaling | 0.807 | ALDH1A2,CYP1B1,FOS |
| Corticotropin Releasing Hormone Signaling | 0.796 | FOS,GLI1,NPR3 |
| Macropinocytosis Signaling | 0.788 | ITGB3,PDGFD |
| Oxidative Ethanol Degradation III | 0.78 | ALDH1A2 |
| DNA damage-induced 14-3-3σ Signaling | 0.78 | SFN |
| Apelin Muscle Signaling Pathway | 0.78 | APLNR |
| Factors Promoting Cardiogenesis in Vertebrates | 0.764 | WNT10A,WNT2,WNT3A |
| Relaxin Signaling | 0.759 | FOS,NPR3,PDE5A |
| The Visual Cycle | 0.759 | RBP2 |
| Valine Degradation I | 0.759 | BCAT1 |
| Fatty Acid α-oxidation | 0.759 | ALDH1A2 |
| Renal Cell Carcinoma Signaling | 0.752 | FOS,SLC2A1 |
| Chemokine Signaling | 0.752 | FOS,Ppp1r12b |
| Role of Macrophages, Fibroblasts and Endothelial | 0.745 | FOS,PDGFD,WNT10A,WNT2,WNT3A |
| Putrescine Degradation III | 0.74 | ALDH1A2 |

| | | |
|---|---|---|
| 3-phosphoinositide Degradation | 0.73 | INPP4A,PPFIA3,PTPRH |
| Xenobiotic Metabolism AHR Signaling Pathway | 0.712 | ALDH1A2,CYP1B1 |
| Cardiac Hypertrophy Signaling | 0.708 | MYL9,RHOJ,RHOV,RND3 |
| Cardiac Hypertrophy Signaling (Enhanced) | 0.708 | EDNRB,ITGB3,LEP,PDE5A,WNT10A,WNT2,WNT3A |
| Ethanol Degradation IV | 0.706 | ALDH1A2 |
| Apelin Cardiac Fibroblast Signaling Pathway | 0.706 | APLNR |
| Breast Cancer Regulation by Stathmin1 | 0.697 | ADGRA2,ADGRD1,APLNR,EDNRB,GPR153,PPP1R14C,PTGER3,PTH1R |
| Insulin Secretion Signaling Pathway | 0.693 | ABCC9,SLC2A1,SLC2A5,SNAP25 |
| Vitamin-C Transport | 0.688 | SLC2A1 |
| Crosstalk between Dendritic Cells and Natural Ki | 0.682 | ACTC1,HLA-A |
| Bupropion Degradation | 0.672 | CYP1B1 |
| Tryptophan Degradation X (Mammalian, via Tryp | 0.672 | ALDH1A2 |
| IL-17A Signaling in Gastric Cells | 0.658 | FOS |
| p53 Signaling | 0.622 | PERP,SFN |
| Sonic Hedgehog Signaling | 0.616 | GLI1 |
| VEGF Signaling | 0.614 | ACTC1,SFN |
| TNFR2 Signaling | 0.604 | FOS |
| Dopamine Degradation | 0.604 | ALDH1A2 |
| Acetone Degradation I (to Methylglyoxal) | 0.59 | CYP1B1 |
| FAK Signaling | 0.585 | ACTC1,ITGB3 |
| Glutathione-mediated Detoxification | 0.578 | HPGDS |
| Ethanol Degradation II | 0.578 | ALDH1A2 |
| T Cell Receptor Signaling | 0.573 | FOS,PTPRH |
| Gαs Signaling | 0.567 | PTH1R,RGS2 |
| CDK5 Signaling | 0.562 | EGR1,PPP1R14C |
| Cytotoxic T Lymphocyte-mediated Apoptosis of T | 0.556 | HLA-A |
| Antioxidant Action of Vitamin C | 0.556 | SLC2A1,SLC2A5 |
| Noradrenaline and Adrenaline Degradation | 0.547 | ALDH1A2 |
| Adrenomedullin signaling pathway | 0.539 | FOS,MYLK,NPR3 |
| B Cell Development | 0.536 | HLA-A |
| IL-17A Signaling in Fibroblasts | 0.536 | FOS |
| Apelin Endothelial Signaling Pathway | 0.524 | APLNR,FOS |
| Neuroprotective Role of THOP1 in Alzheimer's D | 0.52 | HLA-A,Prss30 |
| ERK/MAPK Signaling | 0.52 | FOS,ITGB3,PPP1R14C |
| Inhibition of Matrix Metalloproteases | 0.507 | THBS2 |
| Antigen Presentation Pathway | 0.507 | HLA-A |
| HGF Signaling | 0.5 | FOS,ITGB3 |
| April Mediated Signaling | 0.498 | FOS |
| CD28 Signaling in T Helper Cells | 0.495 | FOS,HLA-A |
| B Cell Activating Factor Signaling | 0.489 | FOS |
| Mechanisms of Viral Exit from Host Cells | 0.489 | ACTC1 |
| CCR3 Signaling in Eosinophils | 0.481 | MYLK,Ppp1r12b |

| | | |
|---|---|---|
| MIF Regulation of Innate Immunity | 0.48 | FOS |
| Estrogen Biosynthesis | 0.48 | CYP1B1 |
| FXR/RXR Activation | 0.472 | FOXA2,SAA1 |
| Serotonin Receptor Signaling | 0.472 | HTR3A |
| IL-6 Signaling | 0.472 | COL1A1,FOS |
| BAG2 Signaling Pathway | 0.472 | ANXA2 |
| 14-3-3-mediated Signaling | 0.467 | FOS,SFN |
| P2Y Purigenic Receptor Signaling Pathway | 0.467 | FOS,ITGB3 |
| tRNA Splicing | 0.463 | PDE5A |
| Retinol Biosynthesis | 0.463 | RBP2 |
| Apelin Pancreas Signaling Pathway | 0.463 | APLNR |
| iNOS Signaling | 0.456 | FOS |
| Gα12/13 Signaling | 0.45 | CDH17,MYL9 |
| Role of Oct4 in Mammalian Embryonic Stem Cel | 0.449 | FOXA2 |
| Sperm Motility | 0.449 | PDGFRA,PDGFRB,PEAK1 |
| Ephrin A Signaling | 0.441 | EFNA3 |
| Adipogenesis pathway | 0.438 | KLF5,LEP |
| MSP-RON Signaling In Cancer Cells Pathway | 0.438 | FOS,SFN |
| Graft-versus-Host Disease Signaling | 0.434 | HLA-A |
| STAT3 Pathway | 0.434 | PDGFRA,PDGFRB |
| Autoimmune Thyroid Disease Signaling | 0.427 | HLA-A |
| Cell Cycle: G2/M DNA Damage Checkpoint Regu | 0.427 | SFN |
| TNFR1 Signaling | 0.42 | FOS |
| Triacylglycerol Degradation | 0.42 | Ces2a |
| Estrogen Receptor Signaling | 0.42 | FOS,LEP,MYL9,Ppp1r12b |
| UVC-Induced MAPK Signaling | 0.413 | FOS |
| D-myo-inositol (1,4,5,6)-Tetrakisphosphate Biosy | 0.407 | PPFIA3,PTPRH |
| D-myo-inositol (3,4,5,6)-tetrakisphosphate Biosyn | 0.407 | PPFIA3,PTPRH |
| UVB-Induced MAPK Signaling | 0.407 | FOS |
| Cardiac β-adrenergic Signaling | 0.407 | PDE5A,PPP1R14C |
| Huntington's Disease Signaling | 0.4 | HIP1,PACSIN1,SNAP25 |
| CD27 Signaling in Lymphocytes | 0.4 | FOS |
| Transcriptional Regulatory Network in Embryonic | 0.395 | GATA6 |
| EGF Signaling | 0.388 | FOS |
| Opioid Signaling Pathway | 0.379 | FOS,GRK5,RGS5 |
| Nicotine Degradation III | 0.377 | CYP1B1 |
| Phagosome Maturation | 0.375 | HLA-A,SNAP25 |
| MSP-RON Signaling Pathway | 0.371 | ACTC1 |
| PKCθ Signaling in T Lymphocytes | 0.362 | FOS,HLA-A |
| Retinoic acid Mediated Apoptosis Signaling | 0.36 | CRABP2 |
| Melatonin Degradation I | 0.36 | CYP1B1 |
| D-myo-inositol-5-phosphate Metabolism | 0.356 | PPFIA3,PTPRH |

| | | |
|---|---|---|
| IL-2 Signaling | 0.355 | FOS |
| Thrombopoietin Signaling | 0.344 | FOS |
| CD40 Signaling | 0.334 | FOS |
| Nicotine Degradation II | 0.334 | CYP1B1 |
| Superpathway of Melatonin Degradation | 0.334 | CYP1B1 |
| Calcium-induced T Lymphocyte Apoptosis | 0.33 | HLA-A |
| Pyridoxal 5'-phosphate Salvage Pathway | 0.33 | GRK5 |
| 3-phosphoinositide Biosynthesis | 0.328 | PPFIA3,PTPRH |
| Serotonin Degradation | 0.325 | ALDH1A2 |
| Remodeling of Epithelial Adherens Junctions | 0.321 | ACTC1 |
| IL-10 Signaling | 0.312 | FOS |
| Erythropoietin Signaling Pathway | 0.309 | FOS,LEP |
| GNRH Signaling | 0.309 | EGR1,FOS |
| Growth Hormone Signaling | 0.307 | FOS |
| T Cell Exhaustion Signaling Pathway | 0.304 | FOS,HLA-A |
| Heparan Sulfate Biosynthesis (Late Stages) | 0.303 | Ces2a |
| T Helper Cell Differentiation | 0.298 | HLA-A |
| Leptin Signaling in Obesity | 0.294 | LEP |
| Regulation of Cellular Mechanics by Calpain Prot | 0.294 | ITGB3 |
| NF-κB Signaling | 0.293 | PDGFRA,PDGFRB |
| Estrogen-Dependent Breast Cancer Signaling | 0.291 | FOS |
| Role of NFAT in Regulation of the Immune Respo | 0.288 | FOS,HLA-A |
| NF-κB Activation by Viruses | 0.287 | ITGB3 |
| GDNF Family Ligand-Receptor Interactions | 0.287 | FOS |
| Toll-like Receptor Signaling | 0.287 | FOS |
| Neurotrophin/TRK Signaling | 0.287 | FOS |
| Antiproliferative Role of Somatostatin Receptor 2 | 0.283 | NPR3 |
| Dopamine Receptor Signaling | 0.283 | PPP1R14C |
| PI3K/AKT Signaling | 0.281 | ITGB3,SFN |
| VDR/RXR Activation | 0.279 | KLF4 |
| IL-7 Signaling Pathway | 0.279 | SLC2A1 |
| IL-3 Signaling | 0.275 | FOS |
| Thyroid Cancer Signaling | 0.275 | FOS |
| Heparan Sulfate Biosynthesis | 0.275 | Ces2a |
| IL-17 Signaling | 0.273 | FOS,LEP |
| JAK/Stat Signaling | 0.272 | FOS |
| Endothelin-1 Signaling | 0.271 | EDNRB,FOS |
| NRF2-mediated Oxidative Stress Response | 0.268 | ACTC1,FOS |
| Prolactin Signaling | 0.268 | FOS |
| Apelin Adipocyte Signaling Pathway | 0.264 | APLNR |
| Xenobiotic Metabolism PXR Signaling Pathway | 0.262 | ALDH1A2,PPP1R14C |
| LPS-stimulated MAPK Signaling | 0.261 | FOS |

| | | |
|---|---|---|
| TR/RXR Activation | 0.257 | SLC2A1 |
| HER-2 Signaling in Breast Cancer | 0.257 | FOS,ITGB3 |
| VEGF Family Ligand-Receptor Interactions | 0.257 | FOS |
| IL-4 Signaling | 0.254 | HLA-A |
| Allograft Rejection Signaling | 0.251 | HLA-A |
| Gap Junction Signaling | 0.249 | ACTC1,NPR3 |
| Superpathway of Inositol Phosphate Compounds | 0.247 | PPFIA3,PTPRH |
| Ceramide Signaling | 0.245 | FOS |
| CTLA4 Signaling in Cytotoxic T Lymphocytes | 0.242 | HLA-A |
| Acute Myeloid Leukemia Signaling | 0.242 | JUP |
| RANK Signaling in Osteoclasts | 0.242 | FOS |
| Altered T Cell and B Cell Signaling in Rheumatoic | 0.238 | HLA-A |
| Regulation of IL-2 Expression in Activated and An | 0.238 | FOS |
| OX40 Signaling Pathway | 0.238 | HLA-A |
| HIF1α Signaling | 0.234 | SLC2A1,SLC2A5 |
| IL-1 Signaling | 0.232 | FOS |
| Death Receptor Signaling | 0.232 | ACTC1 |
| Fcγ Receptor-mediated Phagocytosis in Macroph | 0.227 | ACTC1 |
| CCR5 Signaling in Macrophages | 0.227 | FOS |
| ErbB Signaling | 0.227 | FOS |
| Communication between Innate and Adaptive Imm | 0.221 | HLA-A |
| TGF-β Signaling | 0.221 | FOS |
| Salvage Pathways of Pyrimidine Ribonucleotides | 0.215 | GRK5 |
| UVA-Induced MAPK Signaling | 0.215 | FOS |
| Nitric Oxide Signaling in the Cardiovascular Syst | 0.213 | PDE5A |
| Neuropathic Pain Signaling In Dorsal Horn Neuro | 0.208 | FOS |
| Kinetochore Metaphase Signaling Pathway | 0.208 | PPP1R14C |
| Mouse Embryonic Stem Cell Pluripotency | 0.203 | WNT3A |
| Neuregulin Signaling | 0.198 | ITGB3 |

#### Supplementary Table 2. Sequences of probes for smFISH to detect Zika viral RNA

| Sequence (5' to 3')* |
|---|
| agtctgattcacacagatcaac |
| aaaacctgttgatactgttgct |
| cagaaactctcgtttccaaatc |
| ttagcatattgacaatccggaa |
| ccgtgaatctcaaaaaggctag |
| aacaattccgacactagtatct |
| tccaagtacatatagtatgcac |
| gtatataacacttattcatccc |
| aatcaagtgctttgtgtattct |
| aagtatatgactttttggctcg |
| catgttccaagacaacatcaac |
| agaagccatgtctgatattgat |
| ttcttggagcatgcaaacttag |
| caaatctgaaaagtcaaggcct |
| aggtaatggaatgtcgtggaac |
| tctaagtttatccattttcagg |
| agtacacaaggagtatgacacg |
| cgactcctatgacaatgtaaga |
| ttcaaatgcttttccaatggtg |
| atgatttgaaagctgctccaaa |
| gagaatttgtgagaaccaggac |
| aatagatccattctttgtgttc |
| ggctgtggataagaagatcaac |
| cttcaacgtcgttatagacgaa |
| ttccattcttgaaacagaggag |
| cacgacaaagctgttatttgtc |
| ataatcttctctaaccttgagc |
| cttaacagctgttccaataacg |
| atgttttcatctcgatcagatg |
| agctaaagacttgggtatgatc |
| cattcctcaaaccgaatttcaa |
| gctctgaagatgaaagatacca |
| cttcagagagaggagcataaac |
| aatggtaagttcttcttcacac |
| tgagaccacgtaactgacaatt |
| tacgagtcattactctgtacac |
| catcctttgtcttaaatattcc |
| taacataactcccgtttttgat |
| agtacggagtcttgtttttatg |
| ttgactgctgttgtcatataac |
| cattaagtcgacgatttctgtt |
| tagtagacgtgaagtgaaggtg |
| atacagattatagttggggact |
| ctatacttgagggatctgtgaa |

| Sequence (5' to 3')* |
|---|
| cttgttgaaatgtatcctcttg |
| gaaatgtcagttgtcacgacaa |
| tatgacacggtcagctttaaag |
| atattgtcaaggagcattcttg |

**Supplementary Table 3. Primers used for RT-qPCR**

| Primer | Forward | Reverse |
|---|---|---|
| Human GAPDH | 5’- GCACCACCAACTGCTTAGCAC-3’ | 5’-TCTTCTGGGTGGCAGTGATG-3’ |
| Human IFNε | 5’- AGCACTCATGGGACTGGAAG-3’ | 5’- CAGGTGCTGTAGTCCTGGTT-3’ |
| Murine GAPDH | 5’-TACAGCAACAGGGTGGTGGA-3’ | 5’- GTGAGGGAGATGCTCAGTGT-3’ |
| Murine IFNα | 5’- CCTGAGAGAGAAGAAACACAGCC-3’ | 5’- TCTGCTCTGACCACCTCCCAG-3’ |
| Murine IFNε | 5’-CTCGTCAGTCCAGCAGTGTC-3’ | 5’-TCTGCTGAAGGATCTCGTGC-3’ |
| Murine IFNβ | 5’-AAGAGTTACACTGCCTTTGCCATC-3’ | 5’-CACTGTCTGCTGGTGGAGTTCATC-3’ |
| Murine MX1 | 5’-GAAGGCAAGGTCTTGGATG-3’ | 5’-GCTGACCTCTGCACTTGACT-3’ |
| Murine ISG15 | 5’-GGTACAGAACTGCAGCGAG-3’ | 5’-AGCCAGAACTGGTCTTCGTG-3’ |
| Murine 2’5’-OAS | 5’-CTTTGATGTCCTGGGTCATGT-3’ | 5’-GCTCCGTGAAGCAGGTAGAG-3’ |
| Murine IRF7 | 5’-CCACACCCCCATCTTCGA-3’ | 5’-CCTCCGAGCCCGAAACTC -3’ |
| ZIKV | 5’-TTGGTCATGATACTGCTGATTGC-3’ | 5’-CCTTCCACAAAGTCCCTATTGC-3’ |
